# Selectivity of Protein Interactions along the Aggregation Pathway of α-Synuclein

**DOI:** 10.1101/2021.01.29.428744

**Authors:** André D. G. Leitão, Paulina Rudolffi Soto, Alexandre Chappard, Akshay Bhumkar, Dominic J. B. Hunter, Emma Sierecki, Yann Gambin

## Abstract

The aggregation of α-SYN follows a cascade of oligomeric, prefibrillar and fibrillar forms, culminating in the formation of Lewy Bodies (LB), the pathological hallmarks of Parkinson’s Disease in neurons. Whilst α-synuclein is a major contributor to LB, these dense accumulations of protein aggregates and tangles of fibrils contain over 70 different proteins. However, the potential for interactions between these proteins and the different aggregated species of α-SYN is largely unknown. We hypothesized that the proteins present in the Lewy Bodies are trapped or pulled into the aggregates in a hierarchical manner, by binding at specific stages of the aggregation of α-SYN.

In this study we uncover a map of interactions of a total of 65 proteins, against different species formed by α-SYN. We measured binding to monomeric α-SYN using AlphaScreen, a sensitive nano-bead assay for detection of protein-protein interactions. To access different oligomeric species, we made use of the pathological mutants of α-SYN (A30P, G51D and A53T), which form oligomeric species with distinct properties. Finally, we used bacterially expressed recombinant α-SYN to generate amyloid fibrils and measure interactions with a pool of GFP-tagged potential partners. Binding to oligomers and fibrils was measured by two-color coincidence detection (TCCD) on a single molecule spectroscopy setup. Overall, we demonstrate that LB components are selectively recruited to specific steps in the formation of the LB, explaining their presence in the inclusions. Only a few proteins were found to interact with α-SYN monomers at detectable levels, and only a subset recognizes the oligomeric α-SYN including autophagosomal proteins. We therefore propose a new model for the formation of Lewy Bodies, where selectivity of protein partners at different steps drives the arrangement of these structures, uncovering new ways to modulate aggregation.

**Significance Statement:** The molecular complexity of the Lewy Bodies has been a major hindrance to a bottom-up reconstruction of these inclusions, protein by protein. This work presents an extensive dataset of protein-protein interactions, showing that despite its small size and absence of structure, α-SYN binds to specific partners in the LB, and that there is a clear selectivity of interactions between the different α-SYN species along the self-assembly pathway. We use single-molecule methods to deconvolute number and size of the co-aggregates, to gain detailed information about the mechanisms of interaction. These observations constitute the basis for the elaboration of a global interactome of α-SYN.

## INTRODUCTION

Protein aggregation with formation of intracellular inclusions is a common trait of many neurodegenerative diseases. In Parkinson’s disease (PD) these inclusions are named Lewy Bodies (LB), after the German neurologist Friedrich Lewy who first described them as ‘eosinophilic cytoplasmic bodies, accompanied by neurodegeneration’ in the midbrain^1^. Lewy Bodies are the pathological hallmark of PD and Dementia with Lewy Bodies (DLB) and can also occur in a series of other neurodegenerative disorders including Alzheimer’s disease (AD), Down’s syndrome, Parkinsonism-dementia complex of Guam and neurodegeneration with brain iron accumulation type 1^2^. The key event in the formation of Lewy Bodies is the aggregation of their main constituent, alpha-synuclein (α-SYN), a neuronal protein that is abundantly expressed in the brain^3^. α-SYN aggregates can also be detected in different intraneuronal pathological features: glial cytoplasmic inclusions in Multiple Systems Atrophy (MSA) and axonal spheroids in neuroaxonal dystrophies^4^.

Several factors have been identified as aggregation-triggers of α-SYN. Enhanced oligomerization was first demonstrated by mutations in the α-SYN gene (SNCA) (A30P, E46K, H50Q, G51D and A53T) *in vitro* and *in vivo^5^.* These were the first genetic links to PD to be discovered and occur naturally in early-onset PD patients, who develop Parkinson’s disease earlier in life (in their mid-thirties). Importantly, recent studies revealed that these disease-associated mutants show very different aggregation kinetics, despite causing similar clinical manifestations^6^. The early-onset PD mutants A30P and G51D mainly form oligomers of a smaller size that incorporate WT α-SYN, but do not readily form pre-fibrils; on the other hand, A53T forms larger size oligomers, which do not recruit WT but resemble pre-fibrils^7^.

In 2007, Wakabayashi *et al.^8^* published a comprehensive review that listed all the proteins identified as constituents of Lewy Bodies. Until then, more than 70 proteins had been identified as LB constituents, belonging to 10 different protein classes and involved in a variety of cell functions: structural elements of the LB fibrils; α-SYN-binding proteins; synphilin-1-binding proteins; components of the ubiquitin-proteasome system; proteins implicated in cellular responses such as oxidative and cell stress or chaperones; proteins associated with phosphorylation and signal transduction, cytoskeletal proteins, cell cycle proteins, cytosolic proteins and others.

While many of these proteins are known interactors of α-SYN, the nature of these interactions is still largely unknown. Indeed, as an intrinsically disordered protein (IDP), α-SYN’s function and ability to recruit binding partners is driven by its conformation and degree of oligomerization. The ability to distinguish properties of molecules depending on their self-assembly status is a crucial factor to understand the driving mechanisms of neurodegeneration, where the formation of higher-order assemblies is the norm. With this in mind, we set out to investigate if the proteins present in the Lewy Bodies are trapped or pulled into the aggregates by binding to a particular stage of the aggregation of α-SYN. Most proteins present in the Lewy Bodies are currently assumed as *bona fide* interactors of α-SYN. However, which components of the LB are directly recruited by α-SYN, rather than being recruited by misfolded aggregates of other components within LBs is for most cases unknown. Therefore, a thorough description of the interactome of α-SYN along its aggregation cascade was long overdue.

Here we implemented novel single molecule methods to study co-aggregation and binding events between α-SYN and other proteins present in the LB. To answer the specific challenge of detecting weak and transient interactions, single-molecule fluorescence methods were applied co-translationally to track protein-protein interactions in completely undisturbed samples. We performed measurements directly in the translation reaction of the *Leishmania tarentolae* cell-free protein expression system, bypassing steps of purification or labelling that can modify the oligomerization status of a protein or disrupt interactions.

This work presents strong evidence to the fact that self-assembly of α-SYN dictates its repertoire of binding partners. Strikingly, some proteins in the LB bind to α-SYN with exquisite specificity to different conformers: we defined interactors that bind specifically to monomeric, oligomeric or fibrillar α-SYN, and built a global interactome of α-SYN along the aggregation pathway.

## RESULTS

### Establishing the interactome of α-SYN

The first step in investigating the interactome of α-SYN was to select a list of candidate partners based on our current understanding of Lewy Bodies’ composition. Indeed, many components have been identified in the Lewy Bodies by immunohistochemical methods, and several hundred by mass-spectrometry-based proteomics screens. Here, we chose a subset of 65 Lewy Body components that were grouped by broad functional categories based on the classification established by Wakabayashi *et al^8^.* As expected, these include many components of the proteostasis network such as molecular chaperones, proteins involved in oxidative stress response, aggresome-related proteins, the autophagy-lysosomal pathway and members of the ubiquitin-proteasome system. Structural and signaling proteins are also included (**Figure 1-A** – inset adapted from Brown *et al^9^,* with the author’s consent).

**Figure 1.**
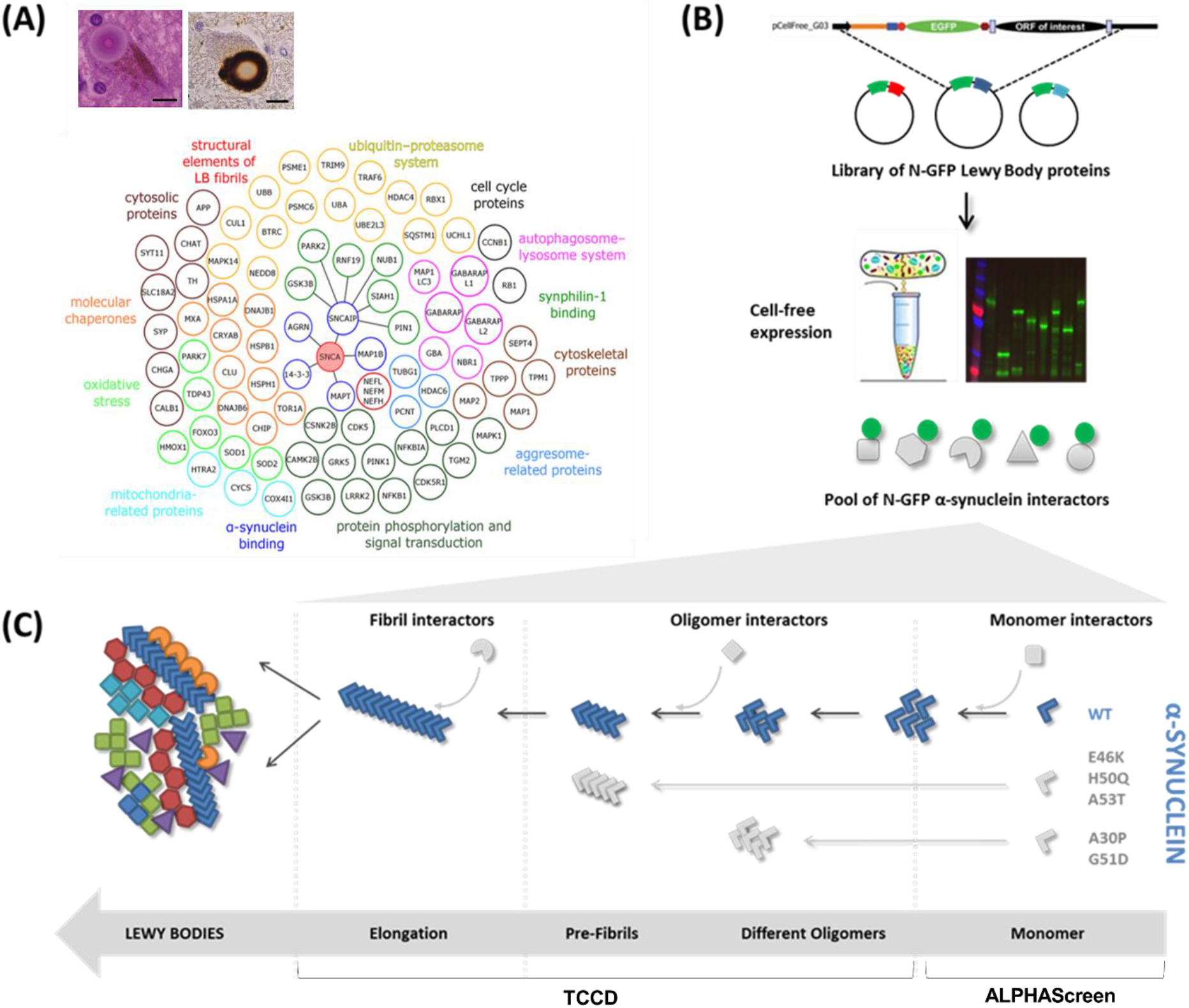
Experimental idea to study the interactome of α-SYN conformers. The goal was to understand the formation of the Lewy Bodies by measuring protein-protein interactions along the aggregation cascade of α-SYN. Figure adapted from Brown *et al^9^* with the author’s agreement. **(A)** The Lewy Bodies contain more than 80 different proteins, which are involved in several cell processes; **(B)** By using a Gateway cloning system, a library of N-terminal GFP tagged Lewy bodies proteins was acquired, for expression in the cell-free *Leishmania tarentolae* lysate; **(C)** On pathway to its inclusion in the Lewy bodies, the aggregation of α-SYN is known for comprising different steps. Upon misfolding of its monomeric form, α-SYN self-assembles into oligomeric, pre-fibrillar and fibrillar forms, respectively. Three different assays were designed to identify the main interactors of α-SYN in four of its forms along this cascade: monomeric (AlphaScreen), oligomeric, pre-fibrillar and fibrillar (two-color coincidence detection using a single-molecule spectroscopy setup).

ORFs encoding these proteins were cloned into the appropriate vectors to allow for expression of N-terminal GFP-tagged proteins using a *Leishmania tarentolae-derived* cell-free expression system (LTE), thus bypassing the need to purify recombinant proteins (**Figure 1-B**). Protein expressions were analysed using SDS-PAGE; resulting bands are depicted in **Figure S1** for a pool of a total of 65 α-SYN potential interactors (see **Table S1** in Annex). All newly cloned N-GFP LB proteins were also sequence-verified, as described in methods section.

Here we focused exclusively on the interactome of α-SYN and to measure protein-protein interactions at the different stages of the aggregation cascade we used a combination of assays: AlphaScreen for monomeric interactions and single molecule fluorescence for oligomeric/fibrillar interactions. The results obtained are described hereafter.

### AlphaScreen reveals LB binding partners of monomeric α-SYN

Direct interactions between monomeric N-terminal mCherry-tagged α-SYN and the N-GFP Lewy Body proteins were assessed using AlphaScreen. AlphaScreen is a proximity assay, where the two interacting proteins bring a “donor” bead and an “acceptor” bead in close proximity. When that happens, the donor beads (coated with the GFP-Lewy Body protein) transfer an excited oxygen singlet to the acceptor beads (coated with mCherry-α-SYN) upon excitation at 680 nm. Reaction with thioxene derivatives in the acceptor bead lead to emission of light at 520-620 nm (**Figure 2-A**). The technique has high sensitivity (up to high micromolar) due to the local increase of the concentration of the proteins at the surface of the beads, allowing the formation of low affinity complexes^10^.

**Figure 2.**
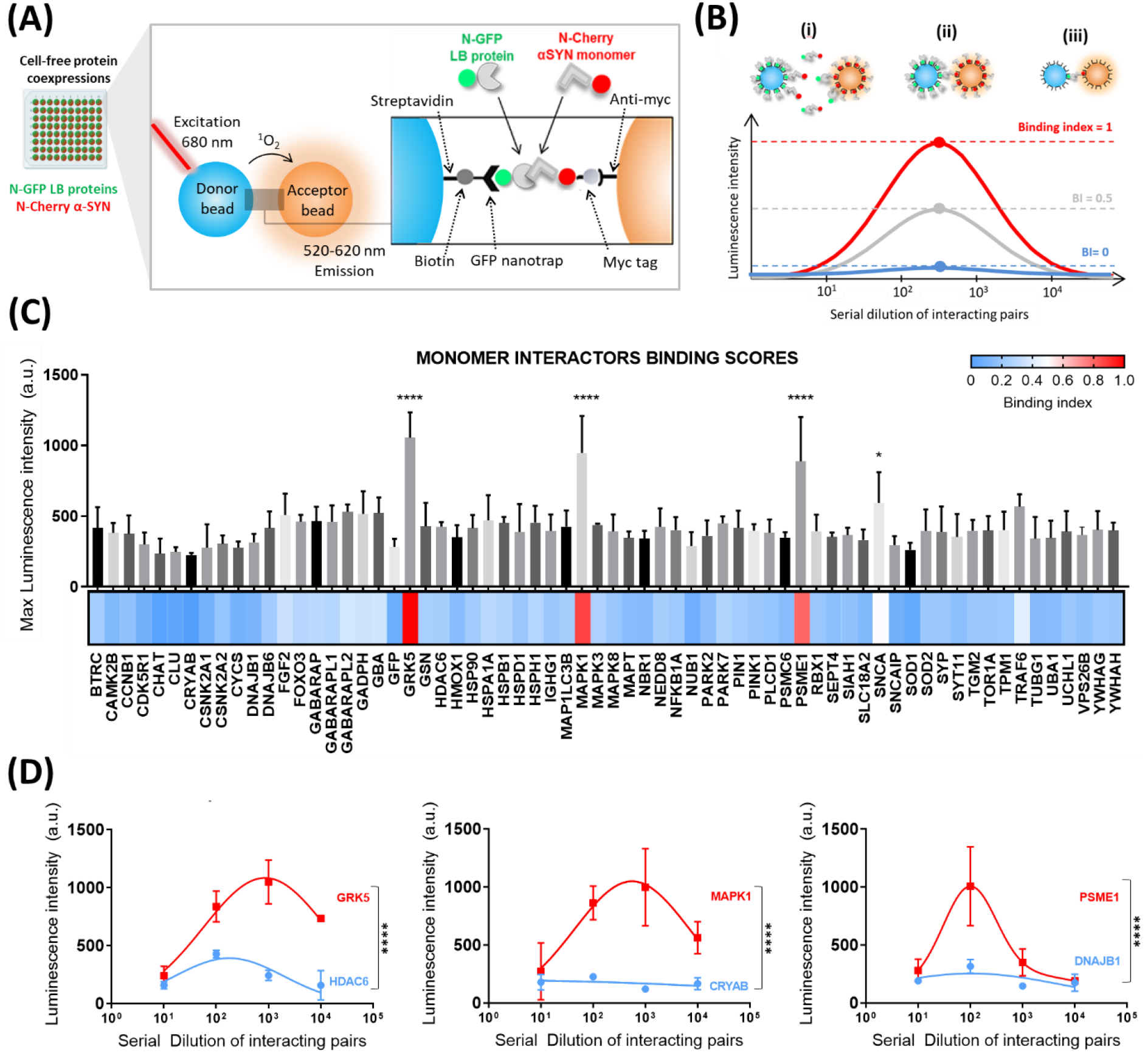
Interactions between LB proteins and monomeric WT α-SYN. **(A)** All Lewy body proteins (see Table S1) and a GFP control were coexpressed in LTE with N-terminal mCherry-tagged WT α-SYN. This was done by priming the LTE with the DNA constructs of each LB protein and α-SYN in a 96-well plate. The donor bead binds the N-terminally GFP-tagged Lewy Body protein while the acceptor bead binds to the N-terminal mCherry-myc tag of α-SYN. When the LB protein and αSYN interact with each other, the proteins will bring the beads in close proximity, with the transfer of a singlet oxygen, leading to AlphaScreen signal being emitted at 520-620 nm. **(B)** AlphaScreen signal is dependent on the dilution of the protein. An excess of proteins **(i)** will lead to a low AlphaScreen signal by inhibition of bead association through competition with the unbound proteins, whereas low concentratrion leads to limited bead association **(iii)**. A maximum signal is detected when the optimal bead/protein ratio is reached **(ii)**. Maximum luminescence is then converted to a binding index (BI) for each pairwise interaction (0<BI<1). **(C)** Maximum values were plotted as a bar plot, with error bars representing the SEM of the triplicate measurements. Heatmap represents the BI values. **(D)** Alphascreen curves along the 4 dilutions revealed 3 main interactors of monomeric α-SYN, resulting in a significant difference in the average maximum signal when compared to that of the GFP control (Dunnett’s multiple comparison’s test; **** p≤0.0001, * p≤0.05, n=3). Error bars = mean ± SEM.

Firstly, mCherry-α-SYN and its GFP protein partners were co-expressed in LTE. Coexpression of the proteins allows for co-translational co-folding to occur, increasing the probability to detect an interaction. Pairwise interactions were then tested by AlphaScreen straight from the cell-free extracts, without any purification steps that could perturb weak complexes. As AlphaScreen signal is dependent on the concentration of the protein (see **Figure 2-B** and **Methods** for a full description of the method), four dilutions were tested for each pairwise experiment in order to observe the expected “hook-effect”. AlphaScreen signals from repeat experiments were then averaged (replicates for several interaction curves are shown in **Figure S2)** and the maxima were normalized to background to give a Binding Index (BI) between 0 (no interaction) and 1 (maximal interaction) (**Figure 2-C**). Most proteins were found to be only weak binders of α-SYN, however, three Lewy Body components emerged as interactors of N-mCherry-tagged α-SYN, when statistically compared to the negative control of monomeric GFP. These interactors include two kinases: G-protein receptor kinase 5 (GRK5) and mitogen-activated protein kinase 1 (MAPK1); and a component of the ubiquitin-proteasome system: proteasome activator complex subunit 1 (PSME1), as illustrated in **Figure 2-D** and maximum AlphaScreen signals plotted in **panel C**. As a control, α-SYN itself was identified as a weak binder, in agreement with our previous data^7^.

### Single Molecule fluorescence reveals selective recognition of LB proteins by oligomeric α-SYN

To rapidly access interactions at the oligomeric level, we performed Two-Color Coincidence Detection (TCCD) experiments. In this assay, both free-floating GFP- and Cherry-tagged proteins are detected by fluorescence confocal spectroscopy. Single Molecule Fluorescence Spectroscopy relies on the detection of fluorescent proteins diffusing in and out of the confocal volume of a microscope (1 femtolitre in volume). In literature, two methods have been predominantly utilised: fluorescence correlation spectroscopy (FCS) and single molecule spectroscopy. While FCS is extremely powerful in correlating diffusion time with size at very small timescales (FCS), it is not optimal when the sample is heterogeneous, as bursts of very large amplitude create long-range temporal correlations. “Pure” single molecule fluorescence techniques are useful when collecting enough events, but are typically performed at pM concentrations, rendering rare events virtually undetectable. Here we used a hybrid approach to overcome some of these limitations by simply analysing the heterogeneity of fluorescence in short time traces, in a background of monomer, performing single particle counting at nanomolar concentrations typically used in FCS experiments.

On the fluorescent time traces acquired in this manner, the diffusion of a large oligomer through the confocal volume results in the appearance of a fluorescent burst well above background. The presence of two such intense fluorescence peaks in both GFP and Cherry channels identifies as a co-assembly event. We utilized a previously described method for automating the scanning of thresholds for each trace, which proved optimal in eliminating noise and detecting a maximum number of events^11^. This method is described by Clarke and colleagues^11^ and was used for all co-expressions in this study, as shown in **Figure S3**.

To access the different oligomeric species, we made use of the pathological mutants of α-SYN, which we reported before to form in the LTE different oligomeric species with distinct properties^7^. We previously observed that A30P and G51D mainly form oligomers of a smaller size that incorporate WT α-SYN, but do not readily form pre-fibrils. On the other hand, A53T forms larger size oligomers, which do not recruit WT but resemble pre-fibrils. Moreover, the inter-molecular FRET was different between the two groups of mutants, indicating differences in conformations^7^. Before carrying out binding assays, the C-terminally mCherry-tagged α-SYN mutants A30P, G51D and A53T were expressed in LTE and observed to behave as described elsewhere for the C-GFP versions of the proteins^7,12^.

To maximize the chances of detecting coincident events, α-SYN mutants and the candidate binding partners were co-expressed (i.e. co-translationally expressed). Furthermore, in order to detect rarer species (that could not be detected when relying solely on Brownian motion) the plate holder of the microscope was adapted to move at a constant set speed during acquisition. In this way we were able to detect sufficient events, even in a background excess of monomer. Using this ‘scanning-plate’ setup, four time traces were acquired for all pairwise co-expressions tested. The binding was here quantified by the Association Quotient ‘Q’ that measures the ratio of coincident events over the total number of events detected, and takes into account the random chances of co-diffusion, by optimising threshold selection as explained in Methods section. Coexpression with monomeric GFP was used as a negative control for this parameter, while coexpression with the N-GFP version of the same protein served as positive control (**panels i and ii in Figure 3-A**, respectively). Here we assume that coexpressing the same forms of synucleins with two different fluorophores provides us with the maximum Q. This maximum was similar for the three mutants tested, with Q~0.5, meaning that, for each selected threshold, approximately half of the protein assemblies detected contained both fluorophores. The other half was assumed either non-coincident or coincident due to chance – chance of co-diffusion is here designated by *E* (**Figure 3-A-iii**).

**Figure 3.**
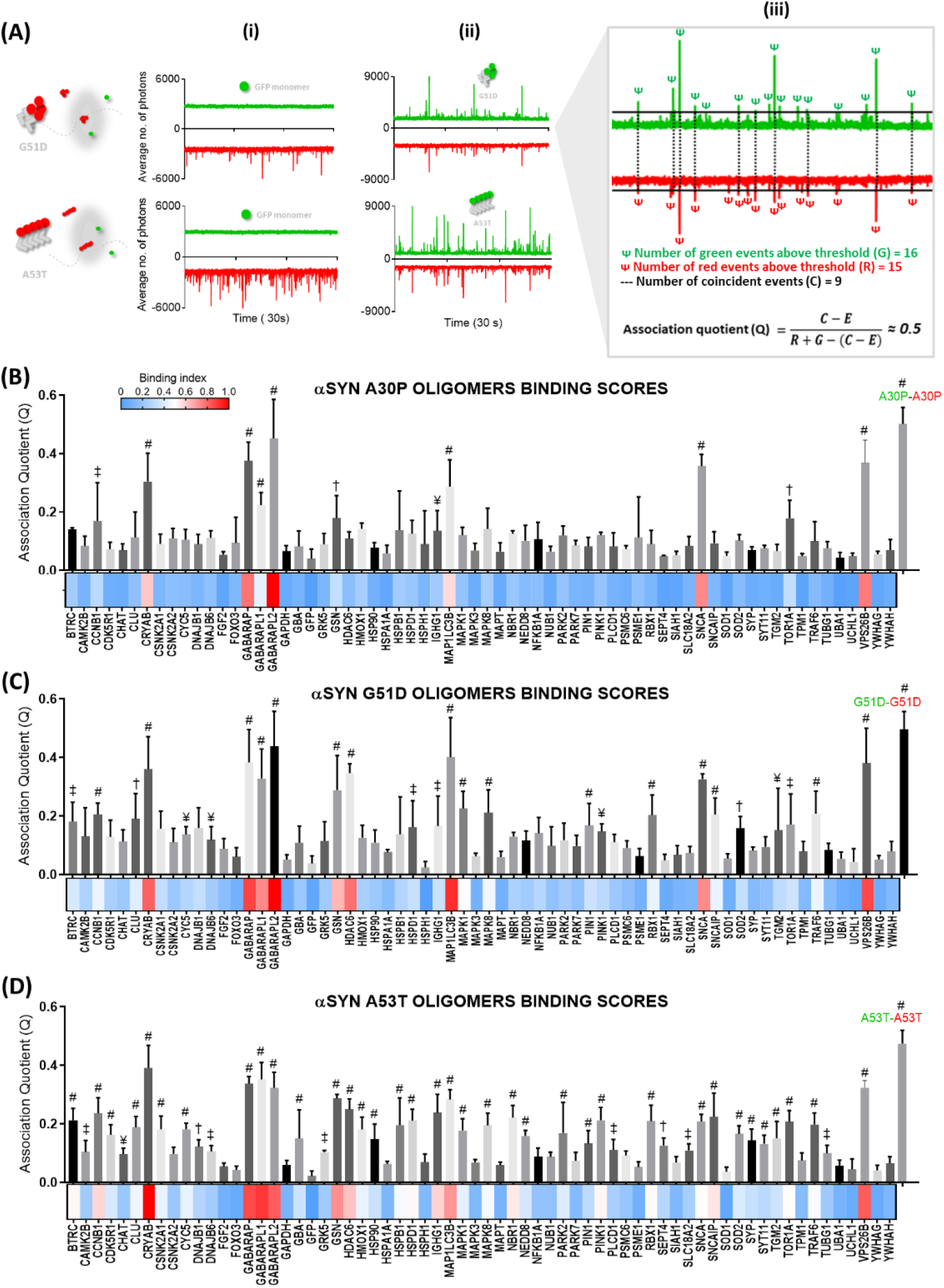
Interactors of oligomeric forms of alpha-synuclein. **(A)** The pathological mutants of α-SYN (A30P, G51D and A53T) were tagged with mCherry in their C-termini and coexpressed in LTE with the library of LB proteins. Shaded circles represent the confocal volume. Four 30 second traces were acquired for each coexpression and TCCD shows binding to synuclein aggregates. **(i)** – A negative control of superfolded GFP was used. **(ii)** – The same mutant form of synuclein, C-GFP tagged, served as a positive control. **(iii)** Association Quotient (Q) was calculated for each trace as the number of coincident events (C) over the total number of events, taking chance coincidence (E) into account. Averages of 4 measurements were acquired and plotted as bar plots and heatmaps for A30P **(B)**, G51D **(C)** and A53T **(D)**. i multiple comparison’s test against coexpressions with GFP. # p≤0.0001, † p≤0.001, ‡ p≤0.01, ¥ p≤0.05. Error bars show mean±SEM. Proteins are displayed in alphabetical order.

The three mutants of α-SYN presented similar interactomes, as shown in **Figure 3-B,C and D**. One of the first observation is that the three main hits from the monomeric interactors do not bind significantly to any of the oligomers. The main interactors of the oligomer species showed average Q values between 0.4-0.5 (similar to the positive control) and did not appear to bind significantly to the monomeric species, already suggesting an interesting selectivity in binding. Interactors of oligomeric species included members of the GABARAP and LC3 (MAP1LC3B) subfamilies, all of which are autophagy-related proteins named ATG8 (traces in **Figure S4**). The small heat-shock protein αβ-crystallin (CRYAB), a molecular chaperone, showed high coincidence values as well (see **Figure S4** for raw trace), whereas other chaperones did not bind to α-SYN oligomers in our assay (see Q values for HSPB1, HSP90, HSPA1A and HSPD1). All the oligomeric species tested here showed high enrichment in the retromer complex subunit VPS26B (vacuolar protein sorting 26 homolog B). In our system, mutant oligomers were not observed to interact with cytoskeleton elements such as tau or tubulin, but the actin-binding protein gelsolin (GSN) was a statistically significant binder for all oligomeric species.

As expected, most of the binding occurs in sub-stoichiometric ratios (meaning that in the co-aggregates, α-SYN is the main component). We measured the number of interactors binding to each oligomer by calculating the ratio of the signals in the two channels. The ratio data for the main binders to A53T aggregates is presented in **Figure S5**. In **Figure S5-A** we show that for the protein concentrations used, apparent stoichiometry did not vary significantly. Our data show that only αβ-crystallin formed complexes of higher stoichiometry with α-SYN. This difference is likely related to the intrinsic propensity of αβ-crystallin to aggregate by itself. Other proteins that co-diffused strongly with α-SYN such as the GABARAPs and MAP1LC3 are predominantly monomeric and showed low presence in heterogeneous aggregates with α-SYN.

### Coaggregation *versus* Binding to pre-formed aggregates

One of the main question is whether the binding occurs as a co-aggregation process during fomation of the oligomers, or whether binding occurs by recognition of a new conformer after the oligomers are fully formed. These two mechanisms can be differentiated by either mixing samples during expression, or mixing samples post-expression. Strikingly, when we performed experiments where interactors of oligomeric α-SYN were mixed after individual expression, few maintained the same Q values (**Figure 4**). Actual recognition of pre-formed and stable α-SYN oligomers seems limited to proteins from the GABARAP/LC3 family that are involved in recognition and enclosure of protein aggregates by autophagosomes during autophagy. Differences in average association quotients (Q values) were statistically significant for all the other identified binders as shown in **Figure 4-B,C**; hence recognition of these proteins occurs during α-SYN oligomerization. This suggests that most of the interactions detected upon coexpression were due to incorporation of the proteins during aggregate formation (as depicted in **Figure 4-B**) although cotranslational co-folding might play a role. Co-aggregators of α-SYN mutants include the small heat-shock protein αβ-crystallin (CRYAB), the retromer complex-associated VPS26B, gelsolin (GSN), HDAC6, as well as synphillin (or syuclein interacting protein SNCAIP) and α-synuclein itself,all of which lose their ability to recognize α-SYN species when experiments are performed in conditions where aggregates are preestablished.

**Figure 4.**
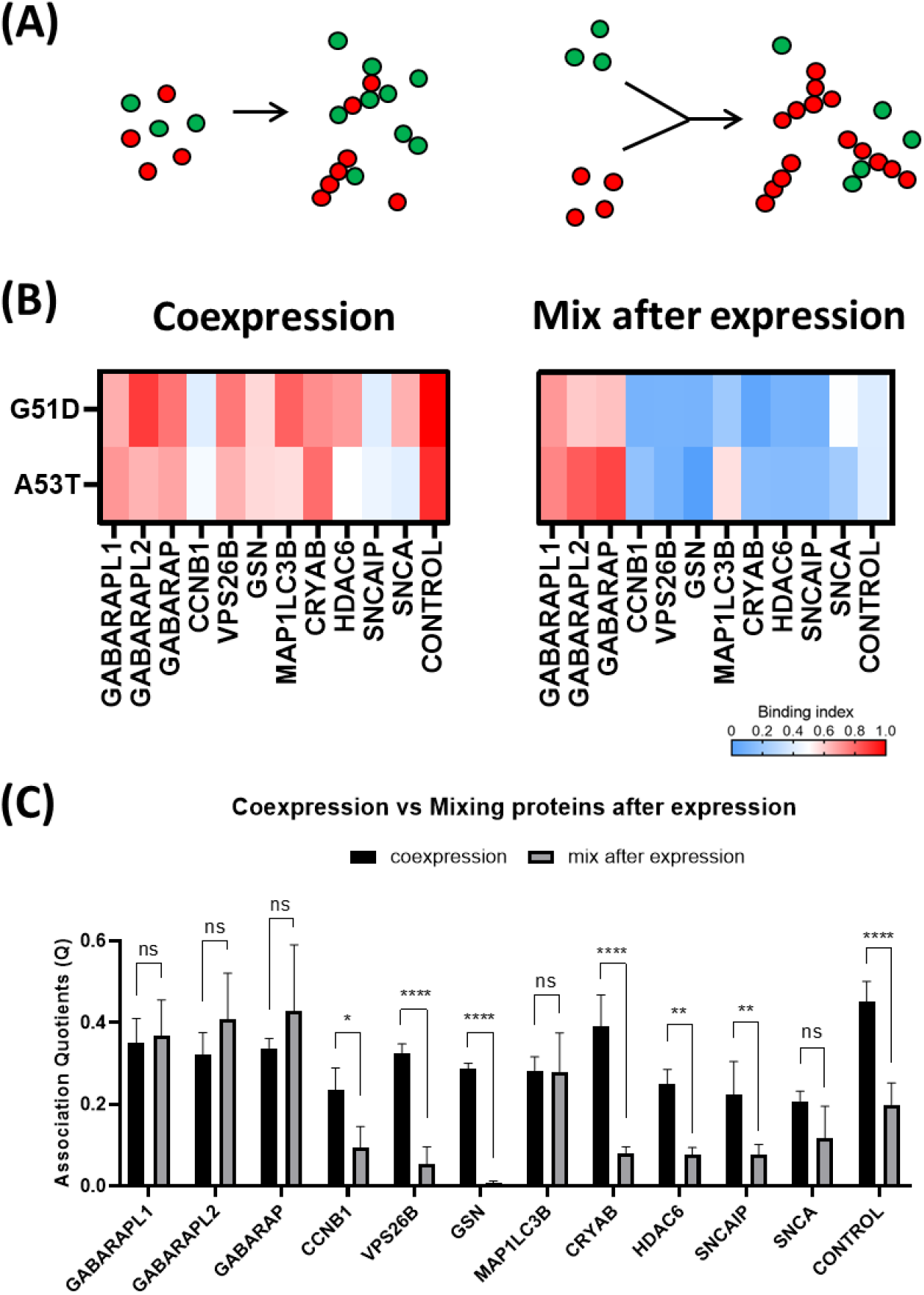
Mixing after expressions reveals binders of pre-formed aggregates. **(A)** Cartoons depicting coexpressing interactors (left) versus mixing interactors with pre-formed aggregates of α-SYN (right). Red/Green circles represent mCherry-tagged synuclein and GFP-tagged Lewy Body component, respectively. **(B)** Heatmap of binding indexes reveals that autophagy-related interactors bind to pre-formed aggregates. Control represents G51D^Cherry^-G51D^GFP^ and A53T^Cherry^-A53T^GFP^ **(C)** Bar plots of Q values with comparison of means using a 2-way ANOVA – Dunnett’s multiple comparison test. **** p≤0.0001, *** p≤0.001, ** p≤0.01, * p≤0.05, n=3. Error bars show mean±SEM.

Note that in physiological conditions, oligomeric species would form in the presence of the various protein interactors, so the co-aggregation processis likely to be relevant. The fact that GABARAP/LC3 proteins have an additional mode of recognition make them strong candidates for furtehr studies.

### Pre-formed fibrils of α-SYN show their own interactome in the LB

We then proceeded to investigate the interactions between LB components and amyloid fibrils of α-SYN. To this end, we used bacterially expressed recombinant α-SYN to generate fibrils, as described in Methods section and illustrated in **Figure 5-A**. These were labelled post-aggregation with the fluorescent dye Alexa594 (in a 1 in 10 ratio) and added to cell-free extracts that were expressing the different binding partners tagged with GFP. Again, TCCD was used to determine binding to the fibrils. In this case, due to the presence of larger and more frequent events (as compared to cell-free expression of mutant synucleins), diffusion due to Brownian motion proved sufficient for detection of enough events, hence a single trace of 300 seconds was acquired for each replicate (**Figure 5-B**) using a stationary plate setup. Overall, we observed a higher apparent binding of Lewy Body proteins, especially many proteins that did not co-diffuse with mutant α-SYN oligomers. Amongst the most striking examples are cytochrome C (CYCS) and chaperones from the DNA-J/Hsp-40 family (here represented by DNAJB1 and DNAJB6), as well as microtubule-associated protein tau (MAPT) (see traces in **Figure 5-D**). Notably, tau was one of the main hits for fibrils and the lowest for oligomeric mutant α-SYN. Such phenomena of specific recognition are addressed in the following results section, in the context of the global interactome of α-SYN. Further, fractions bound to α-SYN fibrils (**Figure S6**) showed considerable differences between interactors; indeed, when expressed at similar levels, interactors such as members of the GABARAP subfamily showed low values of relative presence of GFP species in the aggregates (between ~0.14 for GABARAPL1 and 0.32 for GABARAP), whereas this value was ~0.5 for other hits such as cytochrome C (CYCS). This difference is also noticeable when comparing the traces for coexpression of these proteins in **Figure 5-D**. Again, it should be noted that some of the proteins in the Lewy bodies are intrinsically aggregation-prone, thereby increasing the values of apparent stoichiometry, as could be the case of CDK5R1 and CYCS.

**Figure 5.**
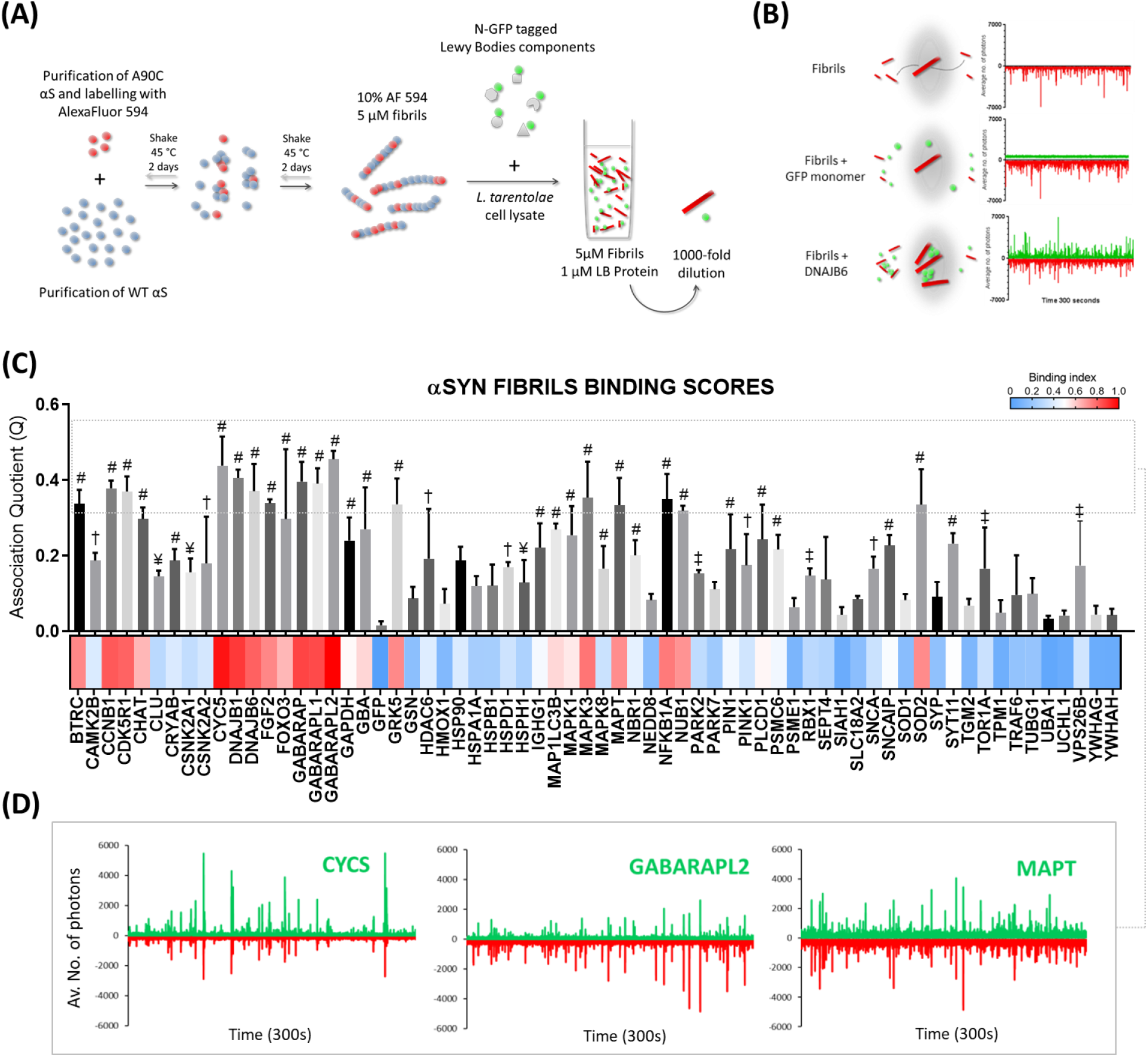
Interactors of pre-formed fibrils of alpha-synuclein. **(A)** Schematic representation of the experimental layout for production of pre-formed α-SYN fibrils (PFF’s). Purified cysteine mutated α-SYN was labelled with AlexaFluor dye 594 (red spheres) and mixed with purified WT α-SYN (blue spheres). After shaking at 45C, fibrils were primed with the cell-free expression reaction of LTE for all the LB proteins tested. After cell-free expression the mixture was diluted 1000 fold to reach single particule concentrations for TCCD experiments. **(B)** Schematic representation of the confocal volumes during measurements of 300s traces. GFP monomer served as a negative control and a known interactor of αSYN as a positive control (DNAJB6) for acquisition of Association Quotients. **(C)** Bar plot of Q values for all the quadruplicated 300s traces and normalized heatmap of binding indexes (BI=0 for GFP monomer and BI=1 for the highest Q value). Error bars represent mean±SEM. Dunnett’s multiple comparison’s test against traces of Fibrils + GFP. # p≤0.0001, † p≤0.001, ‡ p≤0.01, ¥ p≤0.05. **(D)** Traces of 3 of the main interactors, as defined by the top percentile in the distribution of Q values.

An absolute comparison of binding affinities between oligomers and fibrils is difficult. The fact that the apparent number of binders seem higher than for the oligomeric species could be due to the higher concentration of α-SYN aggregates in the assay: fibrils are mixed with interactors at 1 micromolar concentration (monomer equivalent concentration). Also, as the fibrils are larger than oligomers, it becomes easier to detect significant binding as measured in the TCCD experiment. However, as the labelling efficiency of α-SYN amyloid fibrils with AlexaFluor594 was 10%, the real number of binders per α-SYN is therefore lower for fibrils than for oligomers. This make sense as the packing of the amyloid core is very dense and each protein binder would physically occupy a space that would cover multiple α-SYN. Retrospectively, this suggests that binding/co-aggregation with oligomeric species was extremely efficient in our system. In this study, we will therefore focus on the selectivity of interactions observed for each of the aggregated species, rather than on the absolute affinity of interaction(s) between species.

### Concentration effects on Association Quotients

Thus far we have shown that the Association Quotient Q can inform on the binding between Lewy Body proteins and different α-SYN conformers. However, as with any interaction, the concentration of each species (aggregate and binding partner) can affect the overall binding. Although in this study we used a narrow range of protein expressions, such effects should not be overlooked. Therefore, to check that the expression levels of the interactors did not create a bias in the list of binders we plotted average Q values against the expression levels of the GFP-tagged potential partners, for the oligomer- and fibril-binding assays (**Figure S7**). Overall, our data shows that Q can serve as a proxy to identify interactors of synuclein in a high throughput manner, and highlighted important phenomena of selectivity along the aggregation pathway (see **Figure S8** for selective interactors of oligomers – CRYAB – and of fibrils – DNAJB6)

### Global interactome of α-SYN in the Lewy Bodies

The combination of the different datasets defines a global interactome of α-SYN in all its forms (monomeric, oligomeric and fibrillar) *en route* to the formation of Lewy Bodies. The initial pool of interactors was grouped by biological process, according to gene ontology. Information for previously documented interactors was acquired using BIOGRID^13^. Our resulting heatmap of interactions reflects binding indexes (BI), assuming a BI=1 for the most significant interactor. **Figure 6** summarizes this data and highlights specificities in the recognition of the different α-SYN species. Previously reported interactors of α-SYN are also highlighted and further organized in **Figure S10**. Our findings also suggest that not all LB proteins are necessarily interactors of α-SYN and lists newly discovered interactors of α-SYN conformers that had not been identified previously.

**Figure 6.**
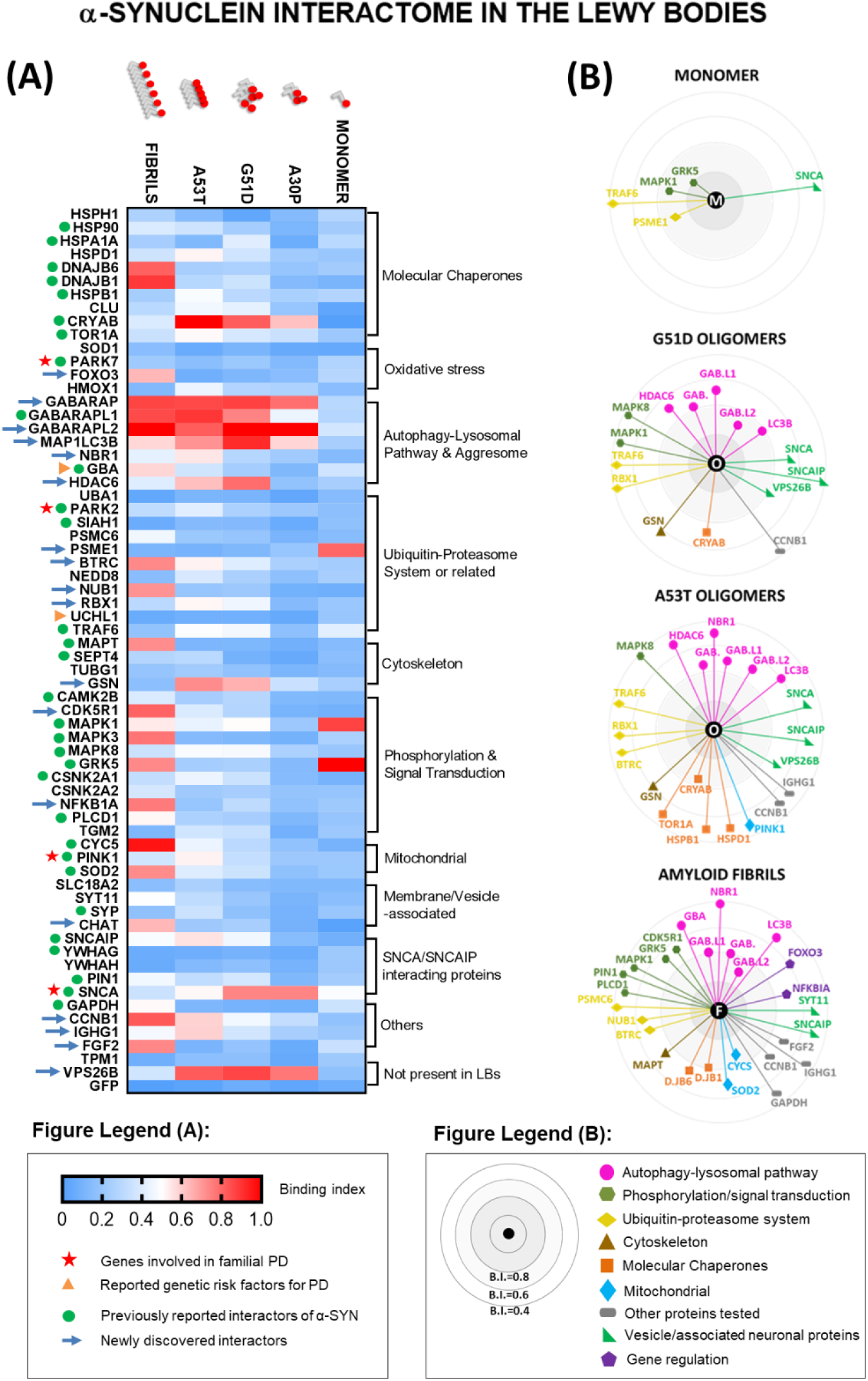
The Interactome of α-Synuclein in the Lewy Bodies. **(A)** Heatmap of interactions reflects binding indexes, distributed from highest (BI=1) to lowest (BI=0), based on Association Quotients *(Q)* from TCCD data and maximum luminescence using Alphascreen proximity assay. Each color depicts the average of triplicate measurements of one pairwise interaction with a LB component. Proteins are organized broadly, based on biological process. Previously identified PPIs (as described in BIOGRID) and genetically relevant proteins are highlighted (see Figure legend). **(B)** Binding indexes allowed for representation of individual interactomes for each species of α-SYN tested (M – monomer, O – G51D/A53T cell-free aggregated species and F – amyloid fibrils). Proteins are grouped by biological process and distance to the center represents change in Binding index.

Strikingly, our single molecule TCCD data reveal a dramatic change in the interactomes, showing that both the oligomeric and the fibrillar state profoundly modify the ability of α-SYN to interact with its environment (**Figure 6**). α-SYN fibrils recruit multiple proteins from the Lewy Bodies; this fits with the fact that α-SYN fibrils are the main component of the LBs and the ones responsible for inclusion formation^14^. More interestingly, our data demonstrate that different aggregated conformers of α-SYN are recognized by different proteins, sometimes with exquisite specificity. This selectivity is further illustrated in **Figure S9**, where we compare exclusivity of binders for each species. A high number of binders showed statistically significant differences in association quotients when comparing fibrils to oligomeric species. Several interactors, including co-chaperones from the DNA-J/Hsp40 family DNAJB1 and DNAJB6, mitochondrial proteins cytochrome C (CYCS) and superoxide dismutase 2 (SOD2), microtubule-associated protein tau (MAPT), kinases such as NF-kappa-β inhibitor alpha (NFKBIA), MAPK3/ERK2 and CDK5R1 (**Figure S9-E**) are specific to fibrillar species. On the other hand, few binders were specific to oligomeric α-SYN: the molecular chaperone αβ-crystallin (CRYAB), gelsolin (GSN), the retromer complex component B (VPS26B) and α-SYN itself (SNCA), which as expected, coaggregated more frequently with early-aggregates than with pre-formed fibrils – **Figure S9-C**. Finally, selective recognition of bigger oligomeric species was detected for histone deacetylase 6 (HDAC6), E3 ubiquitin-protein ligase RBX1 and synphillin (SNCAIP) (**Figure S9-D**), effectively assigning preferential binding of these proteins to different conformations of α-SYN. These phenomena of selective recognition are discussed further below.

## DISCUSSION

### Interdependence between self-assembly and interactome of α-SYN

In this work we demonstrated a biophysical approach to investigate how self-assembly of α-SYN modulates its ability to bind to different partners. Notwithstanding their invaluable contribution to the identification of protein networks, proteomics studies often neglect the fact that for some proteins the formation of supramolecular assemblies is a norm and can profoundly affect partner selection. α-SYN is a good case study, as it is an intrinsically disordered protein (IDP) with multiple binding partners that oligomerizes and self-assembles into higher-order structures.

Overall, our global interactome of α-SYN in the Lewy Bodies uncovers striking phenomena of selectivity along the self-assembly pathway, which could have a profound impact in the progression of disease. Interactors at specific stages could theoretically enhance (i.e. seed) or rescue aggregation (i.e. attempt to inhibit it). As an example of the former, tau and α-SYN are known to synergistically promote fibrillization of each other both *in vitro*^15^ and *in cellulo*^16–17^. Amyloidogenic proteins such as tau and α-SYN share many conformational similarities (notably β-sheet structures), which could explain their cross-seeding. Indeed, we report here that interactions between tau and α-SYN occur predominantly when fibrils of α-SYN are already established. Tau monomers can potentially bind along the surface of α-SYN fibrils, which could then act as a nucleator of tau aggregation, in agreement with other authors^15^.

Other interactors could become trapped in inclusions while attempting to inhibit the aggregation of α-SYN. This could explain why molecular chaperones, proteasomal components and autophagy proteins are ubiquitous in brain inclusions, as discussed below.

### Monomeric interactors of α-SYN

To study the interactome of α-SYN we first used a proximity assay (AlphaScreen). Because of the nature of the assay, AlphaScreen mainly detects interactions between monomers or small oligomers. Perhaps surprisingly, monomeric α-SYN did not bind to the majority of Lewy body proteins tested, including many previously identified interactors of α-SYN. The hit rate of the AlphaScreen experiment is much lower than typically observed in our previous protein-protein interactions screens^10^,^18^,^19^. However, one can also rationalize that the LBs mainly contain interactors of the ‘pathological’ form of α-SYN, while monomers may represent the physiological form of the protein, which could explain the low hit rate observed. As stated before, binding was detected for 2 kinases (G-protein coupled receptor 5 – GRK5 and MAP kinase 1 – MAPK1) and a subunit of the proteasome complex (proteasome activator complex subunit 1 – PSME1) (**Figure 2**).

Previous studies have shown that membrane-bound-GRK5 not only colocalizes with α-SYN in Lewy Bodies^20^, but also phosphorylates Ser-129 of α-SYN at the plasma membrane and induces its translocation to the perikaryal area of neuronal inclusions^21^, consistent with the maturation from pale bodies to Lewy Bodies. Phosphorylation of α-SYN at Ser-129 (pS129-αS) has long been identified as a major modification in LBs^22^. More than 90% α-SYN has been found to be pS129 in the brains of patients with synucleinopathies, as opposed to ~4% phosphorylated α-SYN in healthy brains^22–25^. Importantly, GRK5-catalyzed S-129 phosphorylation also promotes the formation of soluble oligomers and aggregates of alpha-synuclein^20^. Previous studies have shown that α-SYN phosphorylation is mainly occurring at the fibrillary level, after inclusions are formed^26,27^. However, our data suggest that GRK5 phosphorylation could also be an early event, occurring at the monomer level and probably affecting downstream aggregation and/or PPIs.

Other kinases such as MAPK and CDK5 have not been shown to phosphorylate α-SYN^28^, yet they are constituents of LBs^29–32^ and glial cytoplasmic inclusions^33^. Several studies have reported that α-SYN binds MAPK proteins, namely MAPK1 (ERK1)^34^, in agreement with our AlphaScreen findings. It is hypothesized that such interaction reduces the available pool of MAPKS to be phosphorylated by MAPKKs and this is supported by down-regulation of MAPK signalling pathways upon overexpression of α-SYN^34^. Activation of ERK signalling is required for catecholamine^35^ and dopamine release^36^, a crucial step in synaptic transmission.

### Recognition of α-SYN aggregates by Molecular Chaperones

One of the most striking phenomena of selectivity observed in our screen was the different recognition patterns of α-SYN aggregates by molecular chaperones belonging to different families. Here we analysed interactions with several Heat Shock Proteins (HSPs), representative of different groups: Hsp100, Hsp90, Hsp70, Hsp60, Hsp40 and small heat-shock proteins (sHSPs). Such a screen provides valuable information, particularly because the different HSPs exert their function in a concerted action of chaperone networks^37^. In our experimental setup, only three out of the ten HSPs tested were clear hits: the small heat shock protein αβ-crystallin (CRYAB) which showed high Q values for all mutant-generated α-SYN oligomers (and low values for fibrils); and two HSP40/DNAJB family members – DNAJB1 and DNAJB6, which were amongst the main interactors of α-SYN fibrils.

Oligomers have been postulated to be the most toxic species by several studies in the field and several groups have focused on understanding the reason underlying this higher toxicity exerted by oligomers^38,39^. Here we have observed some chaperones recognizing early misfolded states (e.g. co-aggregation of αβ-crystallin with α-SYN), while others were the main hits of PFFs (HSP40s), thus the activity of the two is possibly intimately related. It is noteworthy that none of the other chaperones tested co-diffused with α-SYN forms in our assays. In the case of Hsp70s for example, it is well known that they are not recognizers of protein aggregates *per se,* necessitating an elaborate process of binding and releasing substrates from Hsp40s (co-chaperones)^37,40^.

### Recognition of α-SYN aggregates by Autophagy-lysosomal proteins

Our TCCD data for coexpression of LB proteins with α-SYN oligomers (**Figure 3**) and pre-formed fibrils (**Figure 5**) hint at the importance of autophagy proteins ATG8s (GABARAP and LC3 subfamilies) in the recognition of α-SYN aggregates. Although the structural aspects of this recognition were not studied here, this hints at the involvement of the macroautophagic pathway in the pathogenesis of PD. Crucially, we have also demonstrated that the interaction with ATG8 proteins is due to binding to pre-formed α-SYN aggregates rather than to the incorporation of ATG8s during aggregate formation (**Figure 4**). In sum, ATG8 proteins seem to bind to pre-formed α-SYN aggregates, preferentially in their oligomeric or pre-fibrillar form. One possibility is that such interactions promote packing or clustering of α-SYN conformers and progression towards the formation of inclusions, which could fit with several published reports on the clustering of α-SYN with impairment of macroautophagy^41–43^ (**Figure S11)**.

Autophagy receptors bind to members of the GABARAP/LC3 family through an LC3-interacting region (LIR) that recognizes an LIR-docking site (LDS) in the ATG8 proteins^44^. Interestingly, here we report association between α-SYN aggregates and the same ATG8 proteins without involvement of LIR-containing receptors. Although α-SYN does not show the canonical LIR sequences, the requirements for interactions with ATG8 proteins and the actual function of ATG8s in cargo recruitment remain highly controversial. Structural studies using LIR peptides bound to ATG8 proteins showed that LIRs usually bind as an extended β-sheet to the LDS^44^, and β-sheet formation is a well-described critical step in the aggregation of α-SYN^45–47^. Kalvari and colleagues^48^ have also suggested that binding to ATG proteins is a conformational switching phenomenon (disorder to order transition), which is characteristic of amyloidogenic proteins and could explain our findings.

Although the mechanism by which α-SYN aggregates themselves could recognize GABARAP/LC3 is unclear, the implications for autophagy pathways are interesting. Tanik and colleagues^41^ have demonstrated that seeded α-SYN aggregates are resistant to degradation and impair autophagosome clearance. The authors suggest that α-SYN aggregates interfere with the ability of autophagosomes to mature such that the LC3-positive structures that accumulate in aggregate-bearing cells are incomplete or abnormal and never fuse to lysosomes. Our data fits with these findings and we propose here that binding/clustering of α-SYN aggregates around GABARAP/LC3 could impair their ability to recruit autophagy adaptors for autophagosome growth, transport along microtubules and lysosomal fusion. The fact that GABARAPs and MAP1LC3B do not interact with monomers and do not coaggregate during α-SYN aggregation (instead binding to pre-formed aggregates – **Figure 4**) corroborates this theory.

Autophagy activation by protein aggregates is likely a trade-off between establishing a threshold that is not too low that would lead to exacerbated protein degradation, nor too high that would lead to accumulation of aggregates and rapid impairment of macroautophagy itself. Mutant forms of α-SYN associated with early-onset PD show much higher propensity to aggregate and could destabilize this balance to the latter case, determining much faster disease progressions. Indeed, A30P, G51D and A53T oligomers co-diffused with ATG8s as frequently as mature fibrils (see **Figure 6**), suggesting correlation between these mutant forms and advanced stages of disease. Other authors have shown that α-SYN aggregates also impair autophagy through inhibition of Rab proteins with subsequent mislocalization of ATG9 and defective formation of early vesicles^42,49^. Importantly, restoration of normal autophagic flux reduced ALP impairment and could be an effective strategy in delaying neurodegeneration^50,51^. We propose that understanding the mechanisms of binding between α-SYN aggregates and GABARAPs/LC3 could offer insights on how to restore normal ALP fluxes.

## CONCLUSION

Here we present an extensive dataset to show that despite its small size and absence of structure, α-SYN binds specifically to different partners, and that there is a clear selectivity of protein-protein interactions between the different α-SYN species along the self-assembly pathway. The single-molecule methods used here enable to observe the formation of small co-aggregates when multiple proteins are co-expressed or quantify binding to pre-formed oligomers and mature fibrils. To the best of our knowledge, this is the first study to incorporate the self-assembly of α-SYN into a PPI screen at a large scale, and our data suggest that LB proteins are recruited to the insoluble state at different stages. Overall, this work paints a novel picture of aggregation cascades and protein-protein interactions and lays the groundwork to future studies on how to modulate α-SYN aggregation at different steps.

## MATERIALS AND METHODS

### Preparation of LTE

*Leishmania tarentolae* cell-free lysate was produced as described by Hunter *et al.^52^.* Briefly, *Leishmania tarentolae* Parrot strain was obtained as a LEXSY host P10 from Jena Bioscience GmbH, Jena, Germany and cultured in TBGG medium containing 0.2% v/v Penicillin/Streptomycin (Life Technologies, Carlsbad, CA, USA) and 0.05% w/v Hemin (MP Biomedicals, Seven Hills, NSW, Australia). Cells were harvested by centrifugation at 2500× g, washed twice by resuspension in 45 mM HEPES buffer, pH 7.6, containing 250 mM Sucrose, 100 mM Potassium Acetate and 3 mM Magnesium Acetate and resuspended to 0.25 g cells/g suspension. Cells were placed in a cell disruption vessel (Parr Instruments, Moline, IL, USA) and incubated under 7000 KPa nitrogen for 45 min, and then lysed by rapid release of pressure. The lysate was clarified by sequential centrifugation at 10,000× g and 30,000× g and anti-splice leader DNA leader oligonucleotide was added to 10 μM. The lysate was then desalted into 45 mM HEPES, pH 7.6, containing, 100 mM Potassium Acetate and 3 mM Magnesium Acetate, supplemented with a coupled translation/transcription feeding solution^53^ and snap-frozen until required.

### Gateway cloning to obtain plasmids for cell-free protein expression

We have obtained the Open Reading Frames (ORFs) encoding the desired proteins from IDT. α-SYN point mutants were obtained as gblocks from IDT. A list of Lewy Body components was generated based on a comprehensive review from Wakabayashi and colleagues^8^ (see **Table S1** for all LB proteins used in this study). These genes were sourced from the Human ORFeome collection version 1.1 and 5.1 or the Human Orfeome collaboration OCAA collection (Open Biosystems).

In order to obtain cell-free expression vectors of fluorescently tagged proteins, a gateway cloning method was used, as described elsewhere^54^. Following this protocol, entry clones were generated with PCR primers (Forward primer: 5’GGGGACAAGTTTGTACAAAAAAGCAGGCTT (nnn)_18-25_ 3’, Reverse primer: 5’GGGGACCACTTTGTACAAGAAAGCTGGGTT (nnnn)_18-25_ 3’) – primers to attB1 and attB2 sites respectively (sites flanking the inserts). ORFs were then cloned into Gateway destination vectors designed for cell-free expression^55^: N-terminal GFP tagged (pCellFree_G03), N-terminal mCherry tagged (pCellFree_G05), C-terminal GFP tagged (pCellFree_G04) or C-terminal mCherry-cMyc tagged (pCellFree_G08). The successful clones were selected on LB-Ampicillin (100μg/mL) agar plates, transformed on *E. coli* DH5α competent cells, grown on LB-Amp and the plasmid DNA was extracted using Presto™ DNA miniprep kits and used for cell-free expression with LTE. Following expression, SDS-PAGE gels confirmed the band size (**Figure S1**) and the library of clones was acquired. Finally, all DNA was sequence-verified using Sanger sequencing methods at the Ramaciotti UNSW Center for Cancer and Genomics.

### Alphascreen assay

AlphaScreen is a nano-bead-based proximity assay that allows for rapid and sensitive detection of even weak protein-protein interactions on a cell-free based protein expression system. The method is explained in detail in **Supplementary Appendix I**.

AlphaScreen cMyc detection and Proxiplate-384 Plus 384-wells plates were purchased from Perkin Elmer (MA, USA). Lewy body proteins bearing a N-terminal GFP tag and α-SYN labeled with N-terminal mCherry-Myc were co-expressed in the cell-free system by adding mixed DNA vectors in 10 μL of the *Leishmania tarentolae-based* cell-free system (to a final DNA concentration of 30 nM for the GFP-vector and 60 nM for the Cherry-vector). The mixture was incubated for 3.5 hr at 27°C. Four serial dilutions of the proteins of 1/10 were made in buffer A (25 mM HEPES, 50 mM NaCl). The AlphaScreen Assay was performed in 384-well plates. Per well, 0.4 μg of the Anti-Myc coated Acceptor Beads (PerkinElmer, MA, USA) was added in 12.5 μl reaction buffer B (25 mM HEPES, 50 mM NaCl, 0.001% NP40, 0.001% casein). 2 μl of the diluted proteins and 2 μl of biotin labeled GFP-nanotrap (diluted in reaction buffer A to a concentration of 45 nM) were added to the acceptor beads, followed by incubation for 45 min at room temperature. Then 0.4 μg of Streptavidin coated donor beads diluted in 2 μl buffer A were added, followed by an incubation in the dark for 45 min at room temperature. AlphaScreen signal was recorded on a PE Envision Multilabel Platereader, using the manufacturer’s recommended settings (excitation: 680/30 nm for 0.18 s, emission: 570/100 nm after 37 ms).

Overall, three experiments were done for each pairwise interaction. AlphaScreen signals were averaged to obtain Luminescence curves (**Figure S2**) and the maximum values of those signals were normalized to background to give the Binding Index (BI). Heatmaps of interactions were constructed based on binding indexes (**Figure 2-B-C**).

### Production of pre-formed fibrils (PFFs) of α-SYN

#### Expression and purification of α-SYN for fibril formation

α-SYN WT and A90C (vector pT7-7) were transformed in BL21 (DE3) cells and grown at 37°C in 1L batches of TB medium with 100μg/ml Ampicillin. Cells were induced with IPTG for 4 h and harvested by centrifugation at 9000g, 4°C for 20 min. The supernatant was discarded, and the pellet resuspended in 20 mL lysis buffer (100mM Tris-HCl, 10 mM EDTA, 1x protease inhibitor, pH 8.0) for each Litre grown. The cells were lysed by sonication and the lysate was then boiled for 20 min to denature host protein before centrifugation at 22000g, 4°C for 20 min. The supernatant was collected, and 10 mg/ml streptomycin sulfate was added to remove nucleic acids, followed by another centrifugation at 22000g, 4C for 20 min. The supernatant was again collected, and ammonium sulfate added at a concentration of 0.4 g/mL to precipitate protein. The mixture was stirred for 30 min at 4°C before centrifugation at 22000g. The resulting pellet was resuspended in a minimal volume of 25 mM Tris-HCl pH 7.7 and dialysed overnight against 20 mM Tris pH 7.7. The protein was purified by anion exchange using an HP/Q Sepharose column (GE Healthcare) which was equilibrated with 2 column volumes (120 mL) of wash buffer (20 mM Tris pH 7.7). The sample was injected onto the column using a 50 mL Superloop (GE Healthcare) and eluted over a NaCl gradient from 0 to 2 M collected in 2 mL fractions. SDS-PAGE allowed suitable fractions to be pooled and concentrated using a 2 kDa filter. This was followed by a step size-exclusion chromatography using Superdex G75 column (GE Healthcare) which was equilibrated with 2 column volumes (300mL) of buffer (20 mM NaPO4 pH 7.4). The sample was injected into the column using a 10 mL Superloop and eluted over 1 column volume collected in 2 mL fractions. Suitable fractions were collected, checked on SDS-PAGE and concentrated with a 3kDa filter. Protein concentration was estimated from the absorbance at 275 nm using an extinction coefficient of 5600 M^-1^ cm^-1^ and protein purity was judged by Liquid chromatography-mass spectrometry (LC-MS).

#### Production of AlexaFluor594-tagged α-SYN pre-formed fibrils (PFFs)

Following the protocol developed by Pinotsi and colleagues^56^ (**Figure 5-A**), A90C α-SYN was labelled with maleimide-modified Alexa Fluor 594 dye (ThermoFisher Scientific) via the cysteine thiol moiety. The labelled protein was purified from the excess of free dye by dialysis against PBS at 4 °C overnight, divided into aliquots, flash frozen in liquid N2 and stored at −80 °C.

Pre-formed fibrils were formed by incubating 180 μM of WT α-Syn with 20 μM of A90C α-Syn (final concentration of 200 μM monomeric protein) (PBS, pH 7.4) at 45 °C, stirring with a micro-stirrer. At 24 h intervals, the fibril solution was sonicated using a water bath sonicator for 15 mins. After 72 h, the fibril solutions were divided into 50 μM aliquots, flash frozen with liquid N2 and stored at −80 °C until required. The efficiency of the labelling process was checked by liquid-chromatography mass spectrometry (LC-MS) at the Bioanalytical Mass Spectrometry Facility of the UNSW. This was confirmed by treating the solution of fibrils with an enzymatic detergent at 60°C for 30 minutes to achieve full disaggregation into labelled monomer and by comparing the resulting fluorescence trace with known concentrations of free dye. Labelling efficiency of the α-SYN monomer was determined to be ~10% (i.e. 10% of the monomers were labelled), consistent with Pinotsi *et al*^56^.

For the single molecule experiments, the solutions were diluted to 5 μM (monomer-equivalent) in PBS and sonicated for a further 10 min just before use to obtain a homogeneous distribution of sizes, as described elsewhere^56,57^. Two-color single molecule coincidence experiments were carried out as described above and apparent stoichiometry of interactions with fibrils was calculated.

### Cell-free protein expressions and coexpressions

Briefly, throughout this work 4 types of experiments were performed representing 4 different modalities of cell-free expression of proteins on LTE: a) single protein expressions, either GFP- or mCherry-tagged; b) simple coexpressions of ~1:1 expression levels of proteins tagged with each of the fluorescent tags; c) single protein expressions followed by mixing of the two expression reactions; d) single protein expresssions followed by mixing with purified pre-formed fibrils of α-SYN.

a. For single protein expressions, cell-free expression was carried out by adding DNA to LTE in a ratio of 1:9 and 2:8 for GFP- and mCherry-tagged proteins, respectively. This is because mCherry tags typically display a slower folding time, resulting in overall lower expression levels than GFP-tagged proteins. Proteins were allowed to express for 2.5h at 27°C, followed by 0.5h at 37°C.
b. Coexpressions of GFP and mCherry-tagged proteins were performed in the respective ratios of 20 and 40 nM of DNA template, for a total of 10 μL of LTE. Proteins were allowed to co-express for 2.5h at 27°C, followed by 0.5h at 37°C.
c. Mixing two separate GFP- and mCherry-tagged proteins after individual expression was performed to understand modes of binding for identified interactors. Individual expressions were carried out as above, and mixing was done at 1:1 v/v of the expression reactions after 2h of expression. Mixed samples were allowed to rest at room temperature for 30 minutes before microscope measurements.
d. Finally, we also assessed the binding of N-GFP LB proteins to pre-formed fibrils of α-SYN. Stocks of purified α-SYN fibrils were stored, with 100 μM monomer-equivalent concentration of fibrils. Fibrils were then diluted 1:10 in LTE containing the DNA of LB proteins. Expression was allowed to occur for 2.5h at 27°C, and 0.5h at 37°C, in the presence of fibrils. The reaction was then diluted 1:10 for experiments, to reach a fibril concentration of 1 μM (monomer-equivalent).

All samples acquired as described above were diluted 1:10 in 25 mM HEPES, 50 mM NaCl, directly in the sample holder for microscope measurements.

### Sample preparation and microscope setup

Samples obtained through cell-free expression, as described above, were immediately loaded into a custom-made 192-well silicone plate with a 70 x 80 mm glass coverslip (ProSciTech, Kirwan, QLD, Australia). Plates were analysed at room temperature on Zeiss Axio Observer microscope (Zeiss, Oberkochen, Germany) with a custom-built data acquisition setup. Illumination is provided by a 488 nm and a 561 nm laser beams, co-focused in the sample volume using a 40× magnification, 1.2 Numerical Aperture water immersion objective (Zeiss, Oberkochen, Germany). This creates a very small observation volume in solution (~1 femtolitre), through which fluorescent proteins diffuse, emitting light in specific wavelengths as their fluorescent tags are excited by the laser beams. Light emitted by the fluorophores is split into GFP- and mCherry-channels by a 560 nm dichroic mirror (Dichroic 3). The fluorescence of GFP is measured through a 525/50 nm band pass filter and the fluorescence of mCherry is measured through a long pass filter. Fluorescence is detected by two photon counting detectors (Micro Photon Devices, Bolzano, Italy). Photons of the two channels are recorded simultaneously in 1 milisecond time bins and analysed using LabVIEW 2018 version 18.0 (National Instruments). For experiments with rarer aggregates, in order to increase the efficiency of event detection, the plate holder of the microscope was adapted to move at a constant set speed during acquisition. This step allowed us to retrieve a high number of events, even under excess of monomer.

### Two-color Coincidence Detection (TCCD) to study PPIs

We used the microscope setup described above to excite GFP- and Cherry/AF594-tagged proteins in a cell-free expression system. The identification of fluorescent events in TCCD was achieved by counting the number of photons emitted at a set interval time τ. The presence of two events on both channels at the same bin-time (we used 10ms bin-time) is read as a coincident event.

To perform all our TCCD experiments we followed an optimized methodology published by Clarke and colleagues^11^ to analyze the coexpression traces and measure co-diffusion between dual-labelled molecules, as described next.

#### Association Quotient (Q)

Appropriate thresholds were calculated automatically by plotting the population of Association Quotients (Q) values as a function of the thresholds and finding the maximum Q values. Q is defined as:

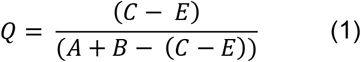

Where *A* and *B* are the events in the two channels, the observed rate of coincident events is C and the estimated rate of events that occur by chance is given by *E*, which in turn can be defined as:

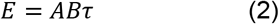

Where *τ* is the interval time in seconds. We used the approach described in the manuscript mentioned above^11^ to calculate, for each time trace of co-expressed m-Cherry- (or AF594-fibrils) and GFP-tagged proteins the optimal threshold, as shown in **Figure S3**.

For each sample, four independent measurements (traces) were acquired and Q values were averaged, to give us an average Q for a specific interaction. We obtained average Q values for coexpressions between N-GFP-tagged LB proteins and C-mCherry α-synucleins. We used the corresponding synuclein C-GFP-tagged as a positive control; coexpressions of C-mCherry synucleins with GFP monomer served as a negative control. For the fibril binding assays we used the maximum Q value for normalization. Average Q values were used as a measure of interactions between protein pairs and binding indexes *(BI)* were calculated by normalizing against the maximum and minimum values detected (positive and negative controls). Binding indexes were used to obtain heatmaps of interactions.

## Acknowledgments

Andre D. G. Leitao was supported by the EMBL Australia Partnership PhD Program. This work was also supported by grants from the National Health and Medical Research Council of Australia (project grants APP1108859 to Yann Gambin and Emma Sierecki, APP1120374 to Emma Sierecki). Yann Gambin was supported by an Australian Research Council Future Fellowship (FT110100478) during this project.

## REFERENCES

1 Holdorff, B. Friedrich Heinrich Lewy (1885-1950) and his work. Journal of the History of the Neurosciences 11, 19–28 (2002).

2 Shults, C. W. Lewy bodies. Proceedings of the National Academy of Sciences 103, 1661–1668 (2006).

3 Spillantini, M. G. et al. α-Synuclein in Lewy bodies. Nature 388, 839 (1997).

4 Dickson, D. W. α-Synuclein and the Lewy body disorders. Current opinion in neurology 14, 423–432 (2001).

5 Conway, K. A. et al. Acceleration of oligomerization, not fibrillization, is a shared property of both α-synuclein mutations linked to early-onset Parkinson’s disease: implications for pathogenesis and therapy. Proceedings of the National Academy of Sciences 97, 571–576 (2000).

6 Li, J., Uversky, V. N. & Fink, A. L. Effect of familial Parkinson’s disease point mutations A30P and A53T on the structural properties, aggregation, and fibrillation of human α-synuclein. Biochemistry 40, 11604–11613 (2001).

7 Sierecki, E. et al. Nanomolar oligomerization and selective co-aggregation of α-synuclein pathogenic mutants revealed by single-molecule fluorescence. Scientific Reports 6, 37630 (2016).

8 Wakabayashi, K., Tanji, K., Mori, F. & Takahashi, H. The Lewy body in Parkinson’s disease: Molecules implicated in the formation and degradation of α-synuclein aggregates. Neuropathology 27, 494–506 (2007).

9 Brown, J. & Horrocks, M. H. in Seminars in Cell & Developmental Biology. (Elsevier).

10 Sierecki, E. et al. Rapid mapping of interactions between human SNX-BAR proteins measured in vitro by AlphaScreen and single-molecule spectroscopy. Molecular & Cellular Proteomics 13, 2233–2245 (2014).

11 Clarke, R. W., Orte, A. & Klenerman, D. Optimized threshold selection for single-molecule two-color fluorescence coincidence spectroscopy. Analytical Chemistry 79, 2771–2777 (2007).

12 Leitao, A., Bhumkar, A., Hunter, D., Gambin, Y. & Sierecki, E. Unveiling a selective mechanism for the inhibition of α-Synuclein aggregation by β-Synuclein. International journal of molecular sciences 19, 334 (2018).

13 Stark, C. et al. BioGRID: a general repository for interaction datasets. Nucleic Acids Research 34, D535–D539 (2006).

14 Froula, J. M. et al. Defining α-synuclein species responsible for Parkinson disease phenotypes in mice. Journal of Biological Chemistry, jbc–RA119 (2019).

15 Giasson, B. I. et al. Initiation and synergistic fibrillization of tau and alpha-synuclein. Science 300, 636–640 (2003).

16 Waxman, E. A. & Giasson, B. I. Induction of intracellular tau aggregation is promoted by α-synuclein seeds and provides novel insights into the hyperphosphorylation of tau. Journal of Neuroscience 31, 7604–7618 (2011).

17 Guo, J. L. et al. Distinct α-synuclein strains differentially promote tau inclusions in neurons. Cell 154, 103–117 (2013).

18 McMahon, K.-A. et al. Identification of intracellular cavin target proteins reveals cavin-PP1alpha interactions regulate apoptosis. Nature communications 10, 1–17 (2019).

19 Moustaqil, M. et al. Homodimerization regulates an endothelial specific signature of the SOX18 transcription factor. Nucleic acids research 46, 11381–11395 (2018).

20 Arawaka, S. et al. The Role of G-Protein-Coupled Receptor Kinase 5 in Pathogenesis of Sporadic Parkinson’s Disease. The Journal of Neuroscience 26, 9227–9238 (2006).

21 Pronin, A. N., Morris, A. J., Surguchov, A. & Benovic, J. L. Synucleins are a novel class of substrates for G protein-coupled receptor kinases. Journal of Biological Chemistry 275, 26515–26522 (2000).

22 Fujiwara, H. et al. α-Synuclein is phosphorylated in synucleinopathy lesions. Nature Cell Biology 4, 160 (2002).

23 Kahle, P. J., Neumann, M., Ozmen, L. & Haass, C. Physiology and Pathophysiology of α-Synuclein: Cell Culture and Transgenic Animal Models Based on a Parkinson’s Disease-associated Protein. Annals of the New York Academy of Sciences 920, 33–41 (2000).

24 Okochi, M. et al. Constitutive phosphorylation of the Parkinson’s disease associated α-synuclein. Journal of Biological Chemistry 275, 390–397 (2000).

25 Anderson, J. P. et al. Phosphorylation of Ser-129 is the dominant pathological modification of α-synuclein in familial and sporadic Lewy body disease. Journal of Biological Chemistry 281, 29739–29752 (2006).

26 Zhou, J. et al. Changes in the solubility and phosphorylation of α-synuclein over the course of Parkinson’s disease. Acta Neuropathologica 121, 695–704 (2011).

27 Walker, D. G. et al. Changes in properties of serine 129 phosphorylated α-synuclein with progression of Lewy-type histopathology in human brains. Experimental Neurology 240, 190–204 (2013).

28 Nakamura, T., Yamashita, H., Takahashi, T. & Nakamura, S. Activated Fyn phosphorylates α-synuclein at tyrosine residue 125. Biochemical and Biophysical Research Communications 280, 1085–1092 (2001).

29 Brion, J.-P. & Couck, A.-M. Cortical and brainstem-type Lewy bodies are immunoreactive for the cyclin-dependent kinase 5. The American Journal of Pathology 147, 1465 (1995).

30 Takahashi, M., Iseki, E. & Kosaka, K. Cyclin-dependent kinase 5 (Cdk5) associated with Lewy bodies in diffuse Lewy body disease. Brain Research 862, 253–256 (2000).

31 Zhu, J.-H., Kulich, S. M., Oury, T. D. & Chu, C. T. Cytoplasmic aggregates of phosphorylated extracellular signal-regulated protein kinases in Lewy body diseases. The American Journal of Pathology 161, 2087–2098 (2002).

32 Kim, E. K. & Choi, E.-J. Pathological roles of MAPK signaling pathways in human diseases. Biochimica et Biophysica Acta (BBA) – Molecular Basis of Disease 1802, 396–405 (2010).

33 Nakamura, S., Kawamoto, Y., Nakano, S., Akiguchi, I. & Kimura, J. Cyclin-dependent kinase 5 and mitogen-activated protein kinase in glial cytoplasmic inclusions in multiple system atrophy. Journal of Neuropathology & Experimental Neurology 57, 690–698 (1998).

34 Iwata, A., Maruyama, M., Kanazawa, I. & Nukina, N. α-Synuclein affects the MAPK pathway and accelerates cell death. Journal of Biological Chemistry 276, 45320–45329 (2001).

35 Cox, M. E. & Parsons, S. J. Roles for protein kinase C and mitogen-activated protein kinase in nicotine-induced secretion from bovine adrenal chromaffin cells. Journal of Neurochemistry 69, 1119–1130 (1997).

36 Bloch-Shilderman, E., Jiang, H., Abu-Raya, S., Linial, M. & Lazarovici, P. Involvement of Extracellular Signal-Regulated Kinase (ERK) in Pardaxin-Induced Dopamine Release from PC12 Cells. Journal of Pharmacology and Experimental Therapeutics 296, 704–711 (2001).

37 Hartl, F. U., Bracher, A. & Hayer-Hartl, M. Molecular chaperones in protein folding and proteostasis. Nature 475, 324–332 (2011).

38 Whiten, D. R. et al. Single-molecule characterization of the interactions between extracellular chaperones and toxic α-synuclein oligomers. Cell reports 23, 3492–3500 (2018).

39 Hinault, M.-P. et al. Stable α-synuclein oligomers strongly inhibit chaperone activity of the Hsp70 system by weak interactions with J-domain co-chaperones. Journal of Biological Chemistry 285, 38173–38182 (2010).

40 Minami, Y., Höhfeld, J., Ohtsuka, K. & Hartl, F.-U. Regulation of the heat-shock protein 70 reaction cycle by the mammalian DnaJ homolog, Hsp40. Journal of Biological Chemistry 271, 19617–19624 (1996).

41 Tanik, S. A., Schultheiss, C. E., Volpicelli-Daley, L. A., Brunden, K. R. & Lee, V. M. Lewy body-like α-synuclein aggregates resist degradation and impair macroautophagy. Journal of Biological Chemistry 288, 15194–15210 (2013).

42 Winslow, A. R. et al. α-Synuclein impairs macroautophagy: implications for Parkinson’s disease. The Journal of Cell Biology 190, 1023–1037 (2010).

43 Yan, J.-Q. et al. Overexpression of human E46K mutant α-synuclein impairs macroautophagy via inactivation of JNK1-Bcl-2 pathway. Molecular neurobiology 50, 685–701 (2014).

44 Birgisdottir, Å. B., Lamark, T. & Johansen, T. The LIR motif-crucial for selective autophagy. J Cell Sci 126, 3237–3247 (2013).

45 Celej, M. S. et al. Alpha-synuclein amyloid oligomers exhibit beta-sheet antiparallel structure as revealed by FTIR spectroscopy. Biophysical Journal 102, 440a–441a (2012).

46 Celej, María S. et al. Toxic prefibrillar α-synuclein amyloid oligomers adopt a distinctive antiparallel β-sheet structure. Biochemical Journal 443, 719–726 (2012).

47 Chen, S. W. et al. Structural characterization of toxic oligomers that are kinetically trapped during α-synuclein fibril formation. Proceedings of the National Academy of Sciences 112, E1994–E2003 (2015).

48 Kalvari, I. et al. iLIR: A web resource for prediction of Atg8-family interacting proteins. Autophagy 10, 913–925 (2014).

49 Gitler, A. D. et al. The Parkinson’s disease protein α-synuclein disrupts cellular Rab homeostasis. Proceedings of the National Academy of Sciences 105, 145–150 (2008).

50 Lee, J.-A. & Gao, F.-B. Inhibition of autophagy induction delays neuronal cell loss caused by dysfunctional ESCRT-III in frontotemporal dementia. Journal of Neuroscience 29, 8506–8511 (2009).

51 Li, L. et al. Parkinson’s disease involves autophagy and abnormal distribution of cathepsin L. Neuroscience Letters 489, 62–67 (2011).

52 Hunter, D. J. B., Bhumkar, A., Giles, N., Sierecki, E. & Gambin, Y. Unexpected instabilities explain batch-to-batch variability in cell-free protein expression systems. Biotechnology and bioengineering 115, 1904–1914 (2018).

53 Johnston, W. & Alexandrov, K. Production of Eukaryotic Cell-Free Lysate from Leishmania tarentolae. Methods in molecular biology (Clifton, N.J.) 1118, 1–15, doi:10.1007/978-1-62703-782-2_1 (2014).

54 Reece-Hoyes, J. S. & Walhout, A. J. M. Gateway recombinational cloning. Cold Spring Harbor Protocols 2018, pdb–top094912 (2018).

55 Gagoski, D. et al. Gateway-compatible vectors for high-throughput protein expression in pro-and eukaryotic cell-free systems. Journal of biotechnology 195, 1–7 (2015).

56 Pinotsi, D. et al. Direct observation of heterogeneous amyloid fibril growth kinetics via two-color super-resolution microscopy. Nano letters 14, 339–345 (2013).

57 Buell, A. K. et al. Solution conditions determine the relative importance of nucleation and growth processes in α-synuclein aggregation. Proceedings of the National Academy of Sciences 111, 7671–7676 (2014).

