## Supporting Information for "Selectivity of Protein Interactions along the Aggregation Pathway of α-Synuclein"

[1]EMBL Australia Node in Single Molecule Science and School of Medical Sciences, The University of New South Wales, Sydney, NSW 2031, Australia. [2]Woolcock Institute of Medical Research, University of Sydney, Sydney, NSW 2037. [33]School of Chemistry, The University of Edinburgh, Edinburgh, EH9 3FJ, United Kingdom. [4]Institute for Molecular Bioscience, The University of Queensland, Brisbane, QLD 4072, Australia.

[+] currently at Department of Molecular Medicine, The Scripps Research Institute, La Jolla, California, CA 92037, United States of America

**SUPPLEMENTARY FIGURE 1:**

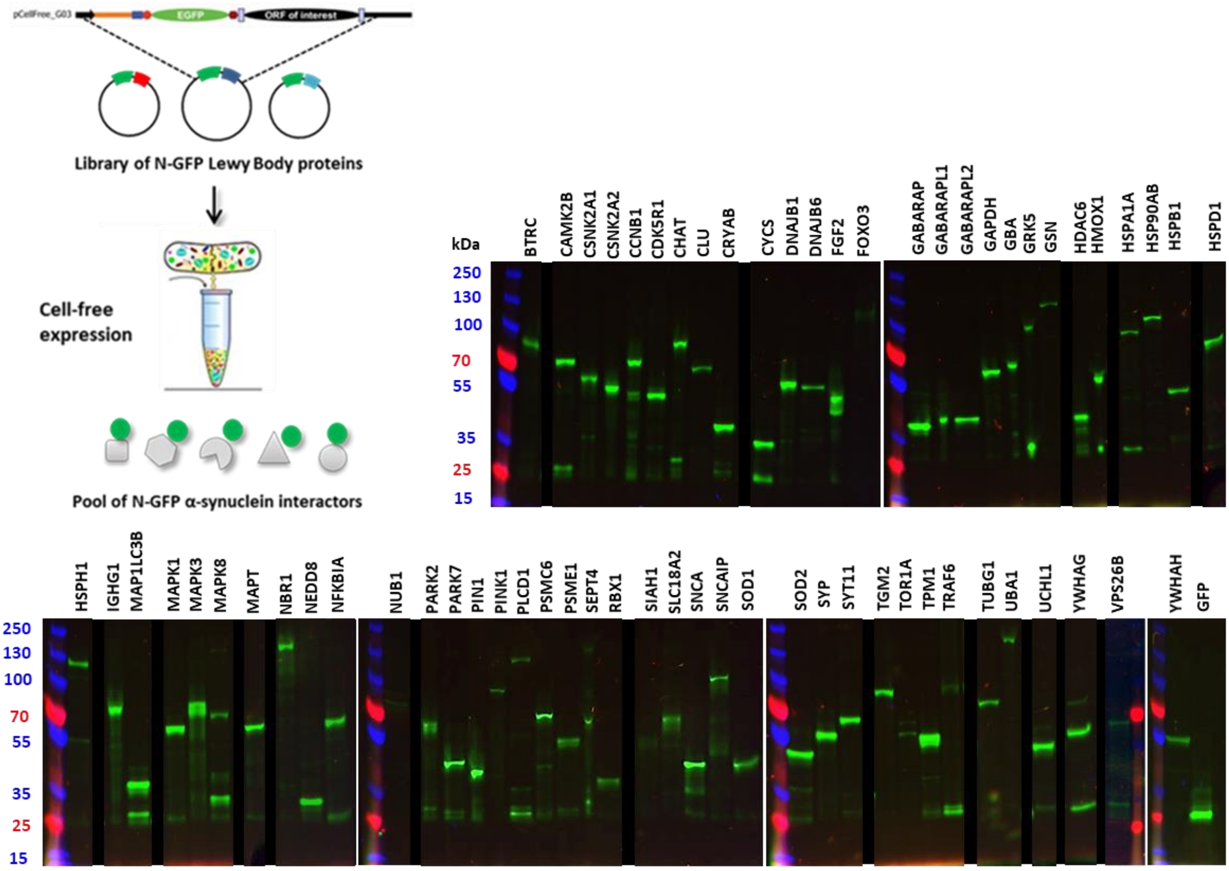


**Figure S1. SDS-PAGE gels of Lewy Body proteins upon cell-free expression.** An library of 65 constructs was cloned using Gateway cloning vectors and obtained DNA constructs were expressed in the cell-free reaction of Leishmania tarentolae extract (LTE). Gels were imaged using ChemiDoc MP imaging system with Image Lab software. Expression was allowed to occur for 2h at 27C and products were loaded on SDS-PAGE gels for quality control of protein expression. Lanes from the same gel were rearranged and are presented here in alphabetical order; rearrangement of lanes is indicated by a black vertical line and a wider space separating each lane. Proteins showing a band at the expected size and with minimal truncation profile were used throughout this study. For full library and sizes of LB proteins see **Table S1**. This supplement figure refers to **Figure 1**.

#### SUPPLEMENTARY TABLE 1:

| GENE | NAME | UNIPROT ID | LENGTH (aa) | MW (Da) | MW + GFP (Da) | Family keyword | Function keyword |
|---|---|---|---|---|---|---|---|
| BTRC | F-box/WD repeat-containing protein 1A | Q9Y297 | 605 | 68,867 | 95,753 | Ubiquitin | Degradation |
| CAMK2B | Calcium/calmodulin-dependent protein kinase type II subunit beta | Q13554 | 666 | 72,678 | 99,564 | Kinase | Brain |
| CCNB1 | G2/mitotic-specific cyclin-B1 | P14635 | 433 | 48,337 | 75,223 | Mitosis | Cell cycle |
| CDK5R1 | Cyclin-dependent kinase 5 activator | Q8N619 | 307 | 34,060 | 60,946 | Kinase | Brain |
| CHAT | Choline O-acetyltransferase | P28329 | 748 | 82,536 | 109,422 | Enzyme | Brain |
| CLU | Clusterin | P10909 | 449 | 52,495 | 79,381 | Chaperone | Folding |
| CRYAB | Alpha-crystallin B chain | P02511 | 175 | 20,159 | 47,045 | Chaperone | Folding |
| CSNK2A1 | Casein kinase II subunit alpha | P68400 | 391 | 45,144 | 72,030 | Kinase | Enzyme |
| CSNK2A2 | Casein kinase II subunit alpha' | P19784 | 350 | 41,213 | 68,099 | Kinase | Enzyme |
| CYC5 | Cytochrome c | P99999 | 105 | 11,749 | 38,635 | Cytochrome | Enzyme |
| DNAJB1 | DnaJ homolog subfamily B member 11 | Q9UBS4 | 358 | 40,514 | 67,400 | Chaperone | Folding |
| DNAJB6 | DnaJ homolog subfamily B member 6 | O75190 | 326 | 36,087 | 62,973 | Chaperone | Folding |
| FGF2 | Fibroblast growth factor 2 | P09038 | 288 | 30,770 | 57,656 | Growth Factor | Apoptosis |
| FOXO3 | Forkhead box protein O3 | O43524 | 673 | 71,277 | 98,163 | Transcription Factor | Gene Regulation |
| GABARAP | GABA(A) receptor-associated protein | Q6IAW1 | 117 | 13,918 | 40,804 | Autophagosome | Degradation |
| GABARAPL1 | Gamma-aminobutyric acid receptor-associated protein-like 1 | Q9H0R8 | 117 | 14,044 | 40,930 | Autophagosome | Degradation |
| GABARAPL2 | Gamma-aminobutyric acid receptor-associated protein-like 2 | P60520 | 117 | 13,667 | 40,553 | Autophagosome | Degradation |
| GAPDH | Glyceraldehyde-3-phosphate dehydrogenase | P04406 | 335 | 36,053 | 62,939 | Enzyme | Gene Regulation |
| GBA | Lysosomal acid glucosylceramidase | P04062 | 536 | 59,716 | 86,602 | Enzyme | Brain |
| GFP | Green fluorescent protein | P42212 | 238 | 26,886 | 53,772 | | |
| GRK5 | G protein-coupled receptor kinase 5 | P34947 | 590 | 67,787 | 94,673 | Kinase | Enzyme |
| GSN | Gelsolin | P06396 | 782 | 85,698 | 112,584 | Actin binding | Brain |
| HDAC6 | HDAC6 protein | Q9BRX7 | 146 | 16,399 | 43,285 | Histone | Gene Regulation |
| HMOX1 | Heme oxygenase | Q96DI8 | 288 | 32,800 | 59,686 | Enzyme | Blood vessel |
| HSP90AB | Heat shock protein HSP 90-beta | P08238 | 724 | 83,264 | 110,150 | Chaperone | Folding |
| HSPA1A | Heat shock 70 kDa protein 1A | P0DMV8 | 641 | 70,052 | 96,938 | Chaperone | Folding |
| HSPB1 | Heat shock protein beta-1 | P04792 | 205 | 22,783 | 49,669 | Chaperone | Folding |
| HSPD1 | 60 kDa heat shock protein, mitochondrial | P10809 | 573 | 61,055 | 87,941 | Chaperone | Folding |
| HSPH1 | Heat shock protein 105 kDa | Q92598 | 858 | 96,865 | 123,751 | Chaperone | Folding |
| IGHG1 | Immunoglobulin heavy constant gamma 1 | P01857 | 330 | 36,106 | 62,992 | glycoprotein | Immunity |
| MAP1LC3B | Microtubule-associated proteins 1A/1B light chain 3B | Q9GZQ8 | 125 | 14,688 | 41,574 | cytoskeleton | Brain |
| MAPK1 | Mitogen-activated protein kinase 1 | P28482 | 360 | 41,390 | 68,276 | kinase | Enzyme |
| MAPK3 | Mitogen-activated protein kinase 3 | P27361 | 379 | 43,136 | 70,022 | kinase | Enzyme |
| MAPK8 | Mitogen-activated protein kinase 8 | A1L4K2 | 427 | 48,088 | 74,974 | kinase | Inflammation |
| MAPT | Microtubule-associated protein tau (amylodogenic isoform 2N4R) | P10636-8 | 758 | 45,850 | 72,736 | aggregate | Brain |
| NBR1 | Next to BRCA1 gene 1 protein | Q14596 | 966 | 107,413 | 134,299 | autophagosome | Brain |
| NEDD8 | neural precursor cell expressed, developmentally down-regulated 8 | Q15843 | 81 | 9,072 | 35,958 | ubiquitin | Degradation |
| NFKB1A | NF-kappa-B inhibitor alpha | P25963 | 317 | 35,609 | 62,495 | transcription factor | Inflammation |
| NUB1 | NEDD8 ultimate buster 1 | Q9Y5A7 | 615 | 70,538 | 97,424 | ubiquitin | Degradation |
| PARK2 | E3 ubiquitin-protein ligase parkin | O60260 | 465 | 51,641 | 78,527 | ubiquitin | Brain |
| PARK7 | Protein/nucleic acid deglycase DJ-1 | Q99497 | 189 | 19,891 | 46,777 | enzyme | Brain |
| PIN1 | Peptidyl-prolyl cis-trans isomerase NIMA-interacting 1 | Q13526 | 163 | 18,243 | 45,129 | isomerase | Enzyme |
| PINK1 | Serine/threonine-protein kinase PINK1, mitochondrial | Q9BXM7 | 581 | 62,769 | 89,655 | kinase | Brain |
| PLCD1 | 1-phosphatidylinositol 4,5-bisphosphate phosphodiesterase delta-1 | P51178 | 756 | 85,665 | 112,551 | lipase | Membrane |
| PSMC6 | 26S proteasome regulatory subunit 10B | P62333 | 389 | 44,173 | 71,059 | proteasome | Degradation |
| PSME1 | Proteasome activator complex subunit 1 | Q06323 | 249 | 28,723 | 55,609 | proteasome | Degradation |
| RBX1 | E3 ubiquitin-protein ligase RBX1 | P62877 | 108 | 12,274 | 39,160 | ubiquitin | Gene regulation |
| SEPT4 | Septin-4 | O43236 | 478 | 55,098 | 81,984 | cytoskeleton | Brain |
| SIAH1 | E3 ubiquitin-protein ligase SIAH1 | Q8IUQ4 | 282 | 31,123 | 58,009 | ubiquitin | Brain |
| SLC18A2 | Synaptic vesicular amine transporter | Q05940 | 514 | 55,713 | 82,599 | transport | Brain |
| SNCA | Alpha-synuclein | P37840 | 140 | 14,460 | 41,346 | membrane association | Brain |
| SNCAIP | Synphilin-1 | Q9Y6H5 | 919 | 100,409 | 127,295 | interacting partner | Brain |
| SOD1 | Superoxide dismutase [Cu-Zn] | P00441 | 154 | 15,936 | 42,822 | enzyme | Brain |
| SOD2 | SOD2 protein | Q96AM7 | 140 | 15,745 | 42,631 | enzyme | Brain |
| SYP | Synaptophysin | P08247 | 313 | 33,845 | 60,731 | membrane association | Brain |
| SYT11 | Synaptotagmin-11 | Q9BT88 | 431 | 48,297 | 75,183 | membrane association | Brain |
| TGM2 | Protein-glutamine gamma-glutamyltransferase 2 | P21980 | 687 | 77,329 | 104,215 | enzyme | Brain |
| TOR1A | Torsin-1A | O14656 | 332 | 37,809 | 64,695 | enzyme | Brain |
| TPM1 | Tropomyosin alpha-1 chain | P09493 | 284 | 32,709 | 59,595 | cytoskeleton | Brain |
| TRAF6 | TNF receptor-associated factor 6 | Q9Y4K3 | 522 | 59,573 | 86,459 | adaptor | Inflammation |
| TUBG1 | Tubulin gamma-1 chain | P23258 | 451 | 51,170 | 78,056 | microtubule | Cytoskeleton |
| UBA1 | Ubiquitin-like modifier-activating enzyme 1 | P22314 | 1,058 | 117,849 | 144,735 | ubiquitin | Degradation |
| UCHL1 | Ubiquitin carboxyl-terminal hydrolase isozyme L1 | P09936 | 223 | 24,824 | 51,710 | ubiquitin | Degradation |
| VPS26B | Vacuolar protein sorting-associated protein 26B | Q4G0F5 | 336 | 39,155 | 66,041 | retromer | Protein transport |
| YWHAG | 14-3-3 protein gamma | P61981 | 247 | 28,303 | 55,189 | scaffold | Signalling hubs |
| YWHAH | 14-3-3 protein eta | Q04917 | 246 | 28,219 | 55,105 | scaffold | Signalling hubs |

**Table S1. LB proteins that were cloned and used for cell-free expression in this study.** Proteins are organized alphabetically from gene name. Length is represented in aminoacids (aa). Molecular weight in daltons (Da). Expected sizes of GFP-tagged constructs are represented in column 'MW+GFP'. Family and function keywords were acquired from Panther Gene Onthology software. This supplementary table refers to **Figure 1** and **Figure S1**.

**SUPPLEMENTARY FIGURE 2:**

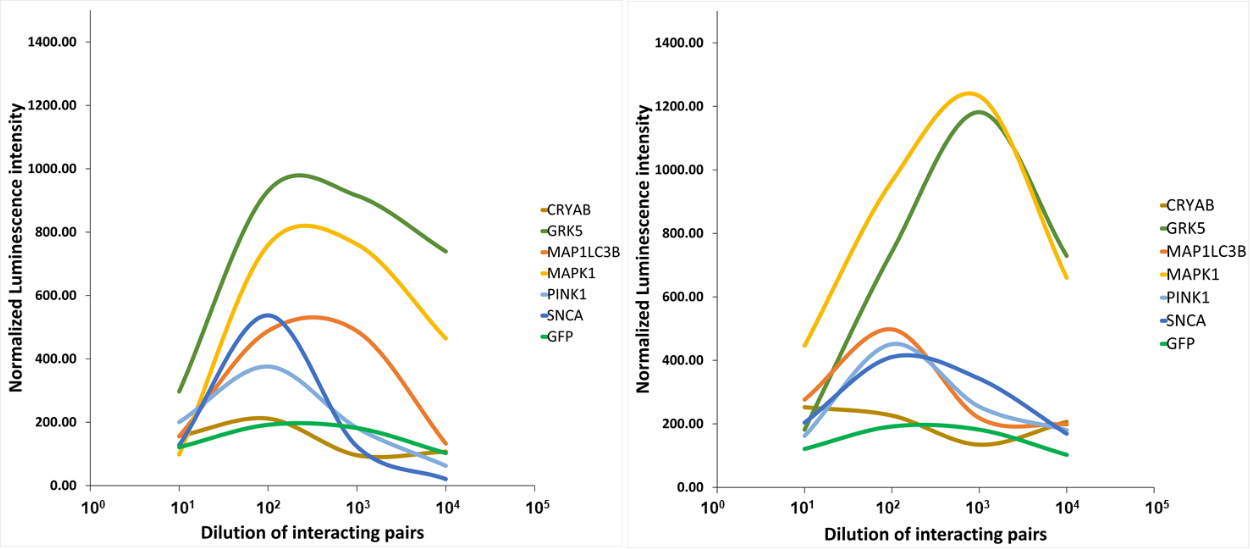


**Figure S2. ALPHAScreen Luminescence curves for cell-free-expressed interacting pairs.** Left and right panels represent two different biological replicates, each depicting average curves of 4 measurements. 7 Different pairwise interactions with α-SYN are shown: 5 GFP-tagged Lewy Body proteins, α-SYN itself and GFP alone. This supplementary figure refers to **Figure 2.**

**SUPPLEMENTARY FIGURE 3:**

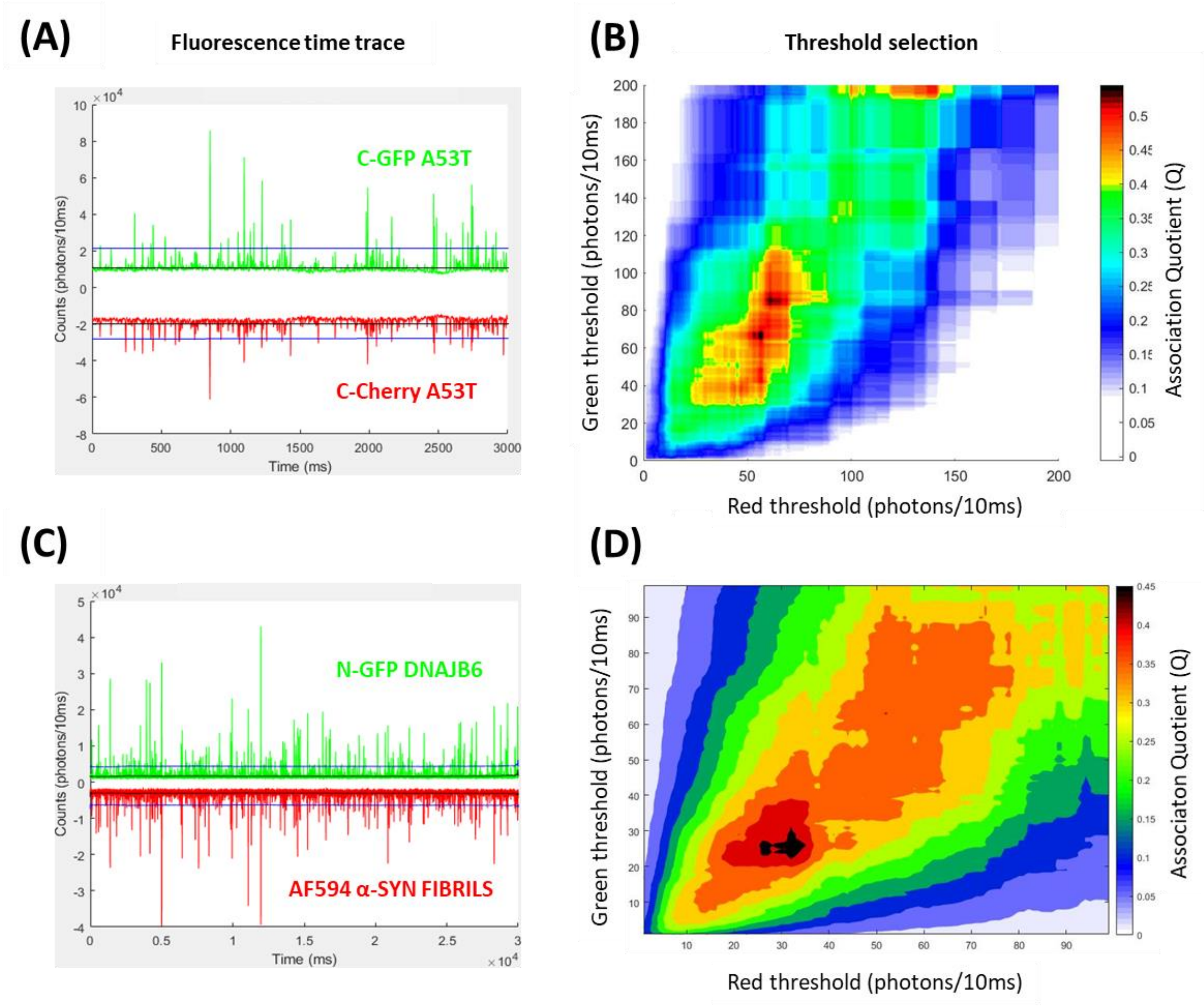


**Figure S3. Optimal threshold selection for calculation of association quotients. (A)** Fluorescence time trace for coexpression of C-GFP A53T and C-Cherry A53T synucleins. Black line (lower) is average intensity; blue line (upper) is the optimal threshold as shown from maximum Q in B. For calculation of threshold and Q, data was binned to 10 ms. **(B)** Color-field map for threshold selection for the trace on the left panel. Optimal thresholds are the ones which give the maximum Q values. Color-field map was acquired by normalizing the binned data from the raw trace to the background. Thresholds were calculated by binning the data to 10ms, in order to filter out small bursts that only cross the threshold for short periods of time (i.e. bursts that cross the threshold for less than 10ms were disregarded). **(C, D)** The same procedure was applied to single-molecule traces of binding to AlexaFluor594-labelled α-SYN fibrils. Data was binned and normalized in a similar manner. This supplementary figure refers to **Figures 3 and 5.**

**SUPPLEMENTARY FIGURE 4:**

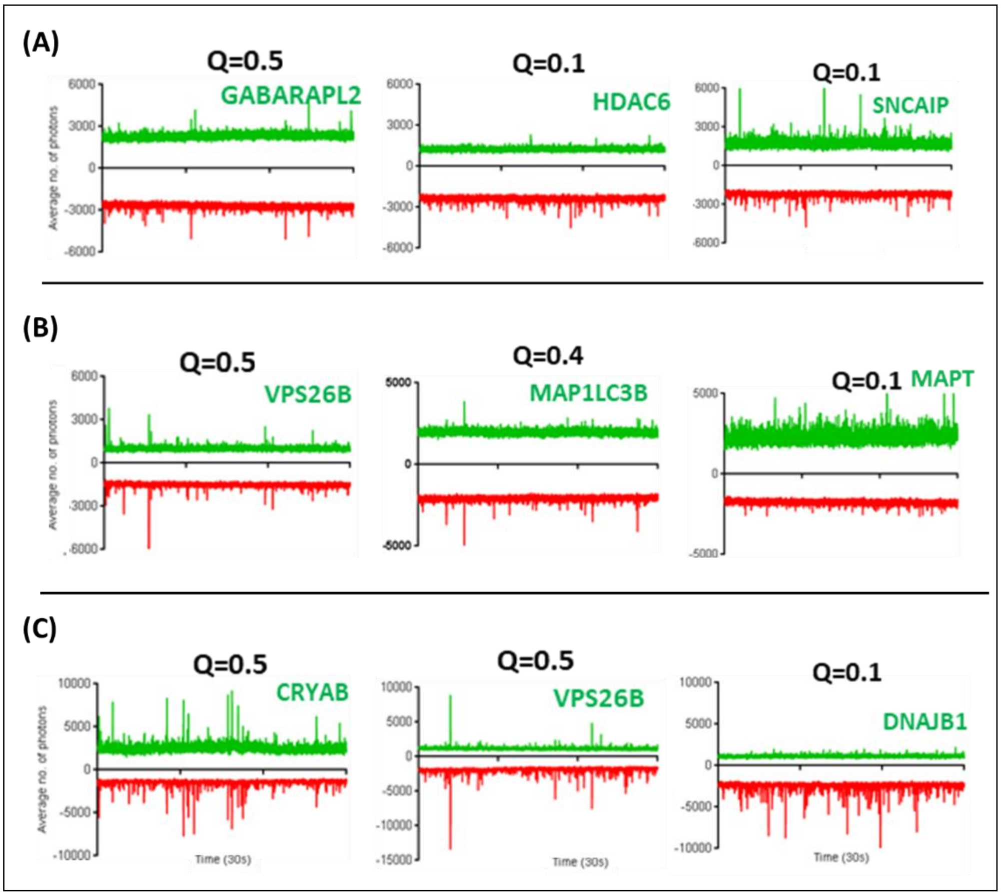


**Figure S4. Example of time traces for cell-free co-expressions between mutant synucleins and Lewy Body proteins.** Fluorescence time trace for coexpression of C-mCherry synuclein and C-GFP Lewy Body proteins: (A, B, C) coexpressions with A30P, G51D and A53T, respectively. Insets represent a single replicate of a 30 second time trace for each co-expression. A total of 4 replicates were obtained and averaged to obtain an average Association Quotient (Q) which was normalized to obtain Binding Indexes throughout this study. This supplementary figure refers to **Figure 3**.

**SUPPLEMENTARY FIGURE 5:**

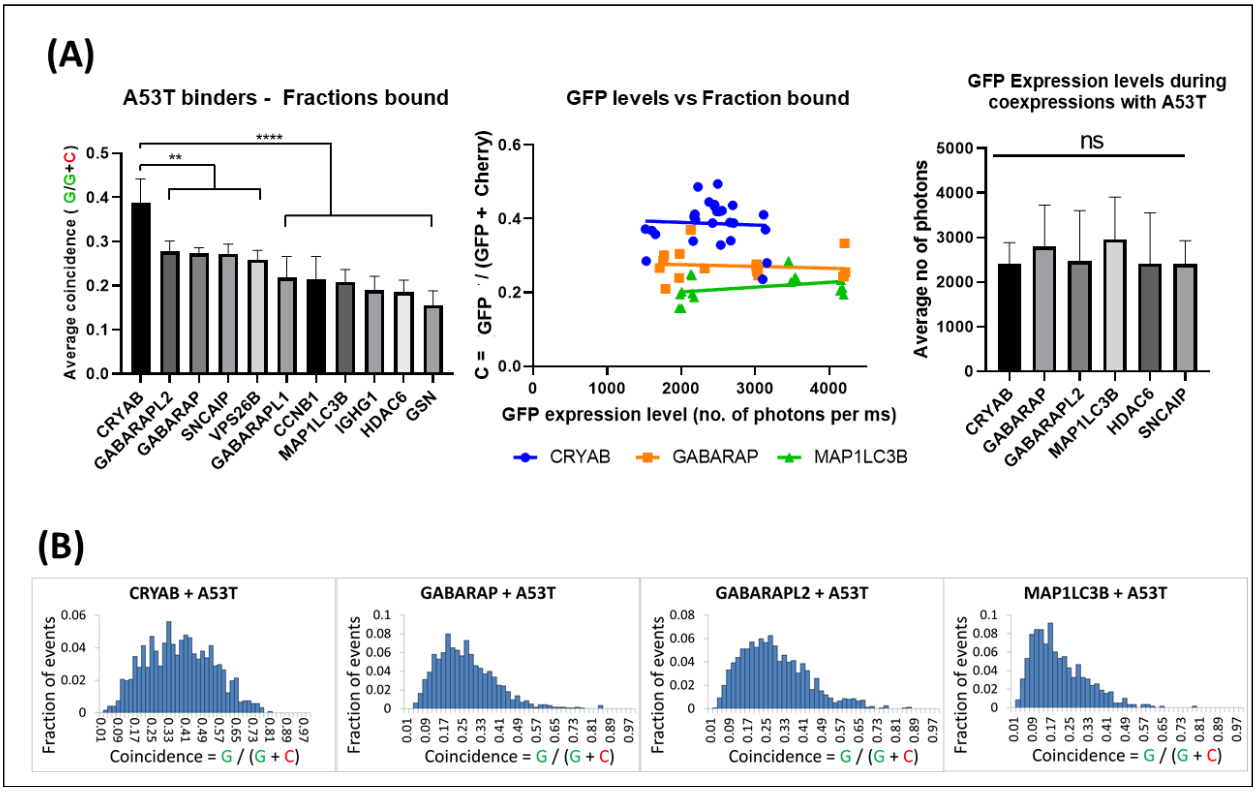


**Figure S5. Analysis of bound fractions for binders of A53T α-SYN conformers. (A)** The variation of GFP expression levels is negligible between each coexpression replicate (middle graph) and all coexpressions were performed at similar GFP expression levels between different proteins (right bar graph). Stoichiometries (left) were measured as the fraction of Cherry-tagged species in coincident events (C=C/(C+G)). **** p≤0.0001, ** p≤0.01 (n=3). **(B)** Distributions of apparent stoichiometries across all the coincident events are depicted. Error bars show mean±SEM. This supplementary figure refers to **Figure 3**.

#### SUPPLEMENTARY FIGURE 6

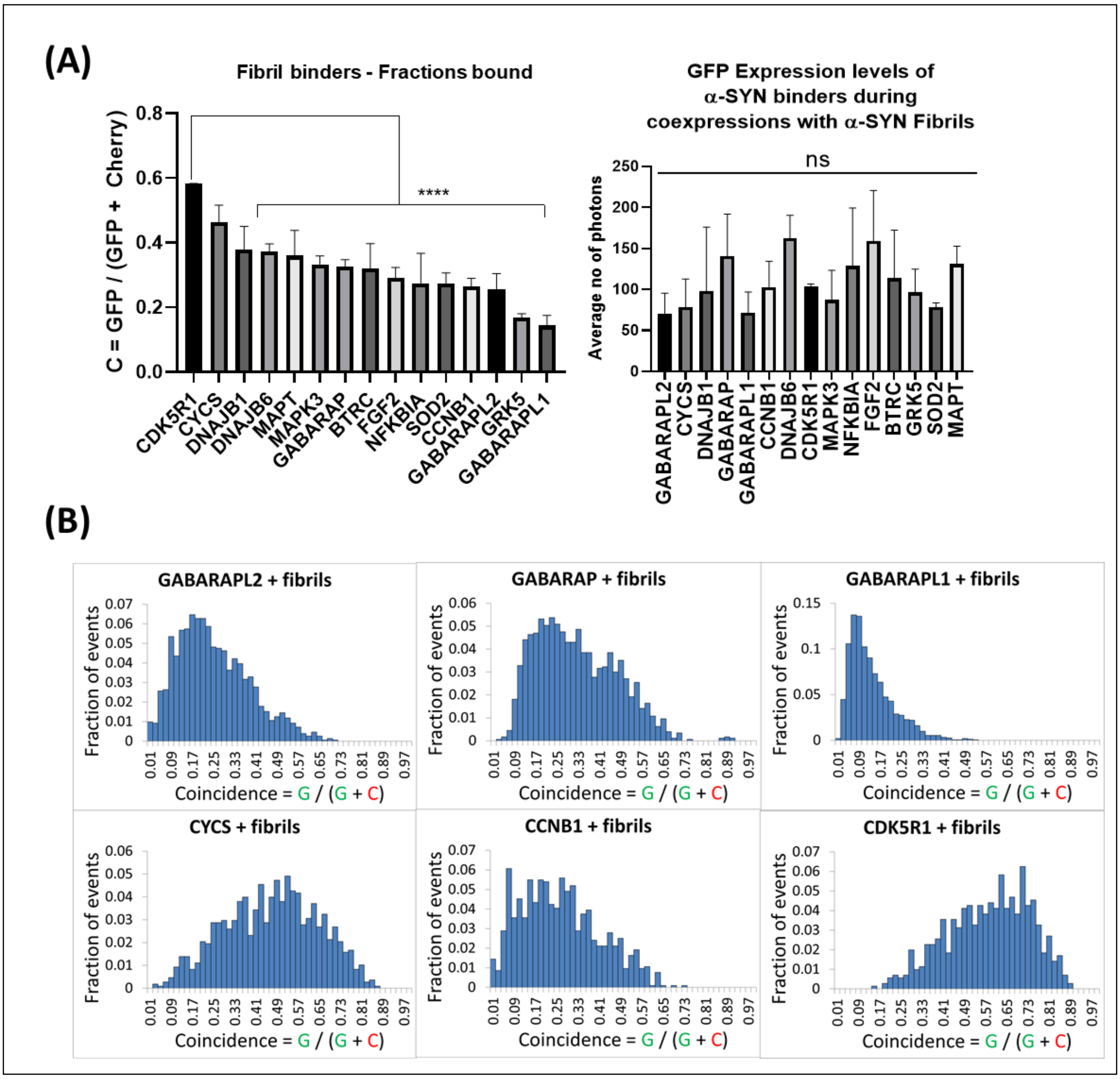


**Figure S6. Analysis of bound fractions for interactions with fibrils of α-SYN. (A)** Differences in bound fractions are shown across the 17 binders with the highest Q values (left graph). GFP expression levels did not vary significantly across all the binders tested (right graph) **(B)** Distributions of coincidence ratios as defined by the relative presence of Cherry species in heterogeneous aggregates. Values represent apparent stoichiometry since these do not account for the 10% incorporation rate of AF594 during labelling. *** p≤0.001, n=3. Error bars show mean±SEM. This supplementary figure refers to **Figure 5**.

#### SUPPLEMENTARY FIGURE 7

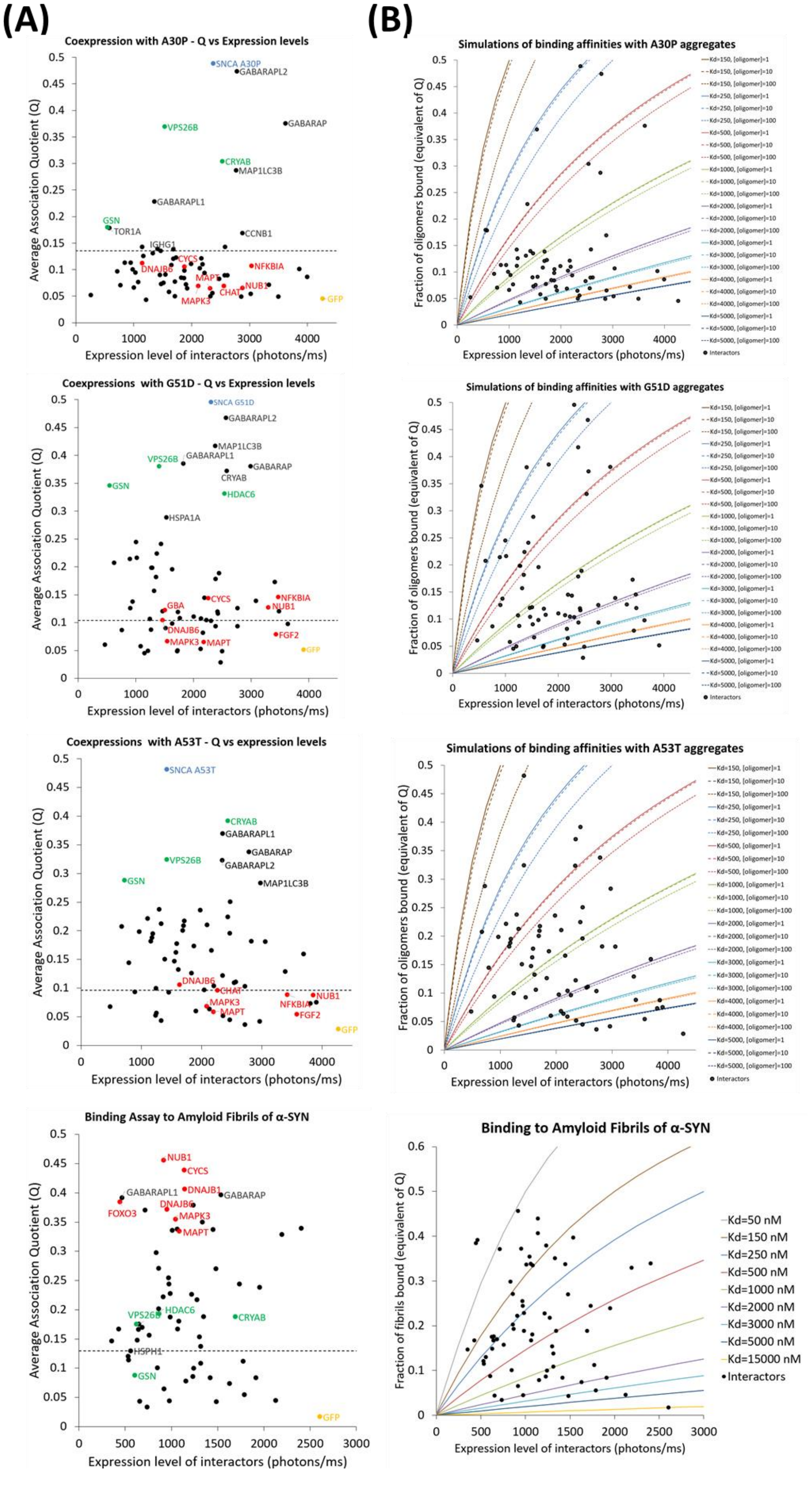

**Figure S7. Effect of concentrations on association quotients with aggregated forms of alpha-synuclein. (A)** Plots of average Q values and expression levels of each interactor. Red – interactors with highest Q values with fibrils; Green – interactors with highest Q values for coexpression with mutants of alpha-synuclein; blue – positive control; yellow – negative control (GFP alone). Dashed line represents the threshold above which Q values are statistically different from Q value of negative control (GFP). **(B)** Plots of dissociation constant (Kd) curves, obtained from simulations using the formula first-order binding Kd. Simulations are plotted for concentration of oligomeric synuclein between 1-100 nM monomer-equivalent. For fibril-binding assay (lower panel), the concentration of fibrils is here corrected for dilution during mixing with GFP-proteins and for the 10% labelling with AF594. Plots show dependence of Q from concentration of interactor and. Sensitivity of the assay between 2000 and 1000 nM of dissociation constants, assuming no second-order binding.

**SUPPLEMENTARY FIGURE 8**

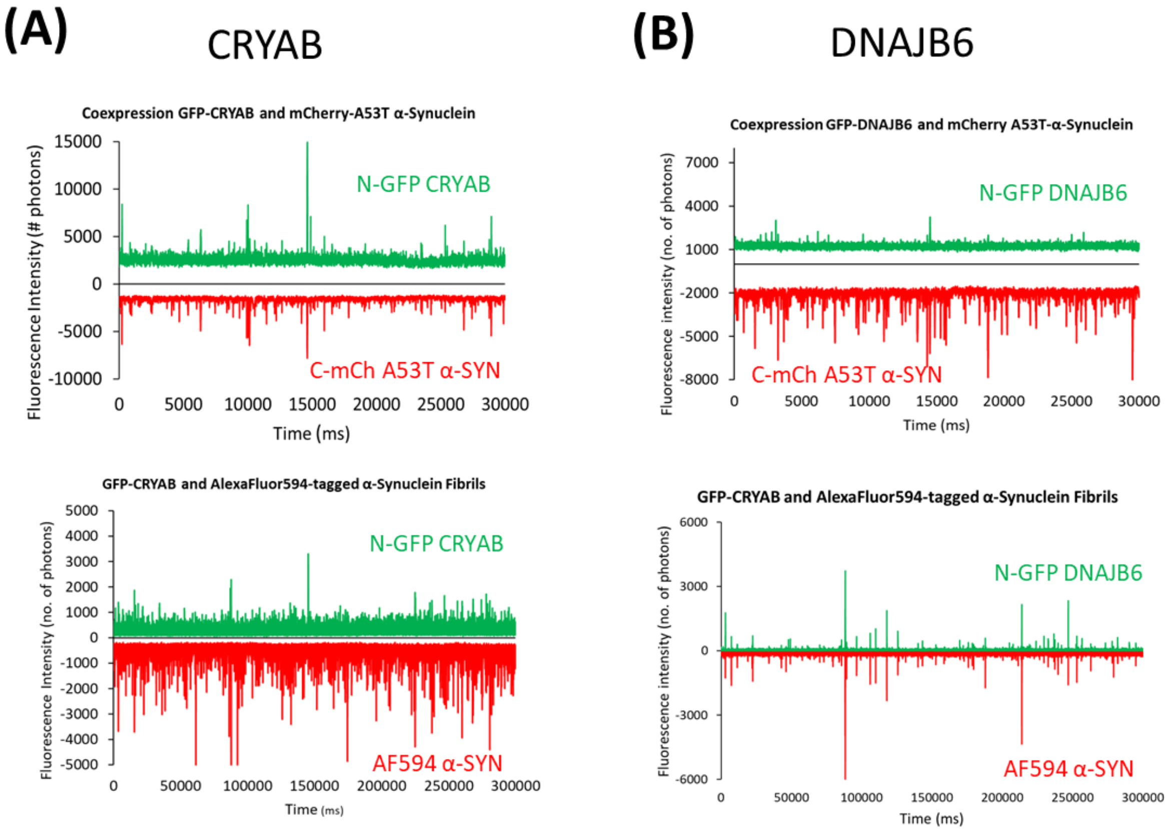


**Figure S8. Selectivity of interactions to A53T oligomers versus amyloid fibrils. (A) Upper panel:** coexpression of the small heat-shock protein alpha-crystallin B (CRYAB) with misfolding mutant A53T, which forms oligomeric structures. Bottom Panel: co-translational expression of CRYAB in cell-free extract with purified amyloid fibrils of alpha-synuclein.(B) Similar setup for the HSP40 protein, DNAJB6 showing no binding to oligomeric A53T and binding to fibrils.

**SUPPLEMENTARY FIGURE 9**

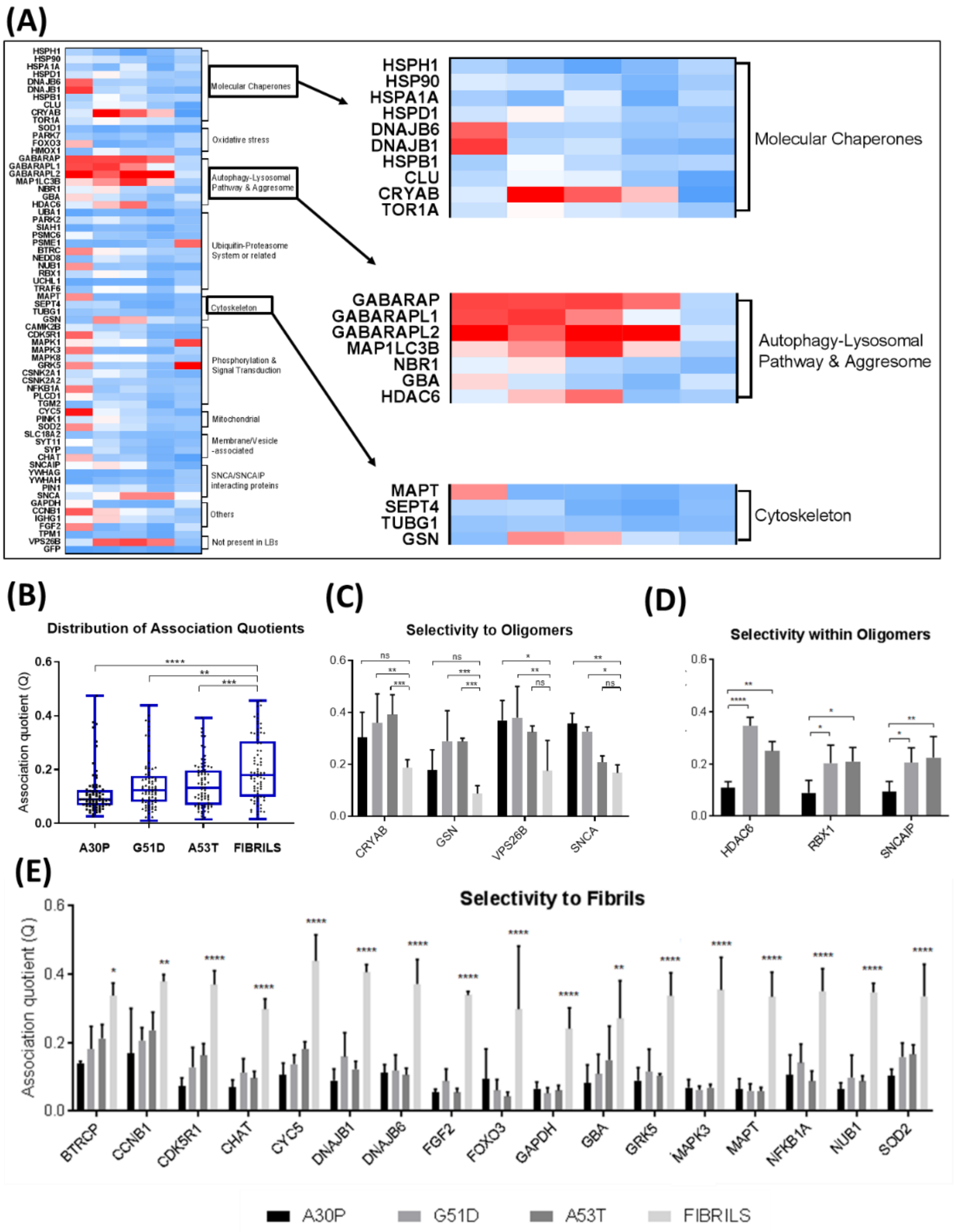


**Figure S9. Selectivity in the recognition of α-SYN aggregates. (A)** Heatmap of interactions highlighting specificities within protein families. **(B)** Q values were distributed for all the synuclein forms tested. **(C)** Selectivity to oligomers was defined as a statistically significant difference between one or more mutant forms of cell-free expressed synuclein and PFFs. **(D)** Few proteins showed a higher Q value for one of the mutant forms, as compared to the other two (selectivity within oligomers). **(E)** Selectivity to fibrils was defined as a statistically significant higher Q value for binding to fibrils as compared to the highest oligomer Q value of the same pairwise interaction. (Dunnett's multiple comparison test, **** p≤0.0001, *** p≤0.001, ** p≤0.01, * p≤0.05, n=3). Error bars show mean±SEM. This supplementary figure refers to **Figure 6**.

**SUPPLEMENTARY FIGURE 10**

**(A)**

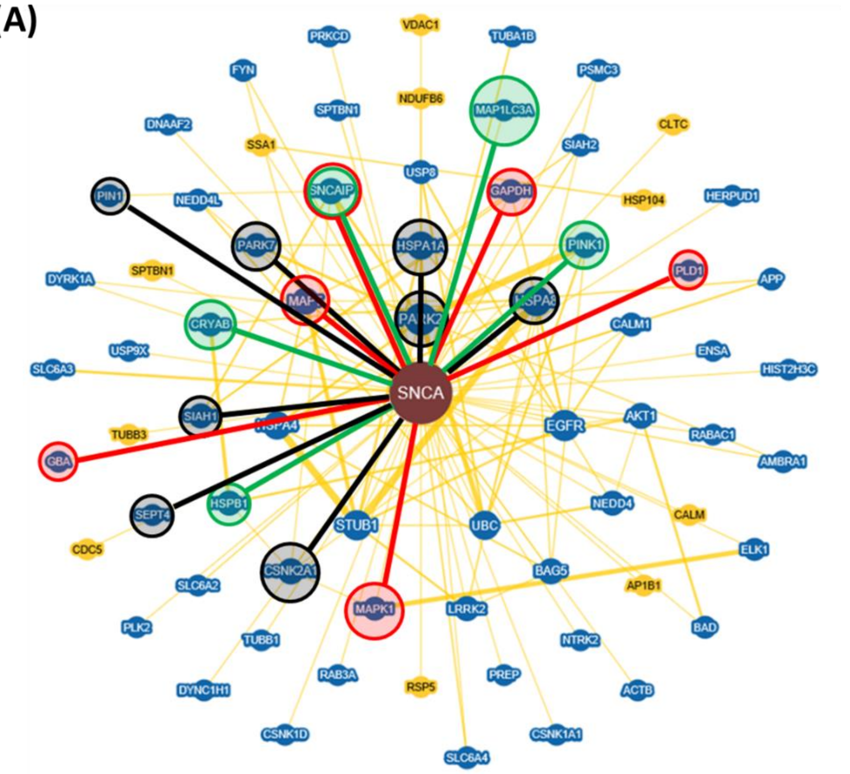


**(B)**

| Previously documented interactors validated | Newly discovered interactions | Documented Interactors not validated | Interaction not documented and not Detected | Present in LBs but not tested | |
|---|---|---|---|---|---|
| | | | | SNCA interactors | No PPI evidence |
| DNAJB1 | FOXO3 | HSP90 | HSPH1 | APP | CHIP |
| DNAJB6 | GABARAP | HSPA1A | HSPD1 | MAP1B | PCNT |
| CRYAB | GABARAPL2 | HSPB1 | CLU | LRRK2 | CHGA |
| TOR1A | MAP1LC3B | PARK7 | HMOX1 | SQSTM1 | AGRN |
| GABARAPL1 | NBR1 | PARK2 | UBA1 | MAP1S | HDAC4 |
| GBA | HDAC6 | SIAH1 | PSMC6 | MAP2 | |
| MAPT | PSME1 | TRAF6 | NEDD8 | | |
| MAPK1 | BTRC | SEPT4 | TUBG1 | | |
| MAPK3 | NUB1 | CAMK2B | CSNK2A2 | | |
| GRK5 | GSN | MAPK8 | TGM2 | | |
| PLCD1 | CDK5R1 | CSNK2A1 | SLC18A2 | | |
| CYCS | NFKBIA | SYP | SYT11 | | |
| PINK1 | CHAT | YWHAG | YWHAH | | |
| SOD2 | CCNB1 | PIN1 | TPM1 | | |
| SNCAIP | IGHG1 | | GFP | | |
| SNCA | FGF2 | | | | |
| GAPDH | VPS26B | | | | |
| | RBX1 | | | | |

**Figure S10. Alpha-synuclein interactome in the LBs (comparison with published data). (A)** Interactome of α-SYN, based on list of peer-reviewed published interactors, available in BIOGRID. **(B)** Interpretative table for the interactome of α-SYN. Highlighted interactions in insets A and B: red – interacted preferentially with fibrillar α; green – oligomeric mutant α-SYN (A30P, G51D or A53T); blue – monomeric WT α-SYN; black – no interaction was detected in this study; grey – previously described as present in LBs but not tested in this study - the high molecular weight of these proteins made them unsuitable for cell-free expression in *Leishmania tarentolae*. This supplementary figure refers to **Figure 6**.

#### SUPPLEMENTARY FIGURE 11

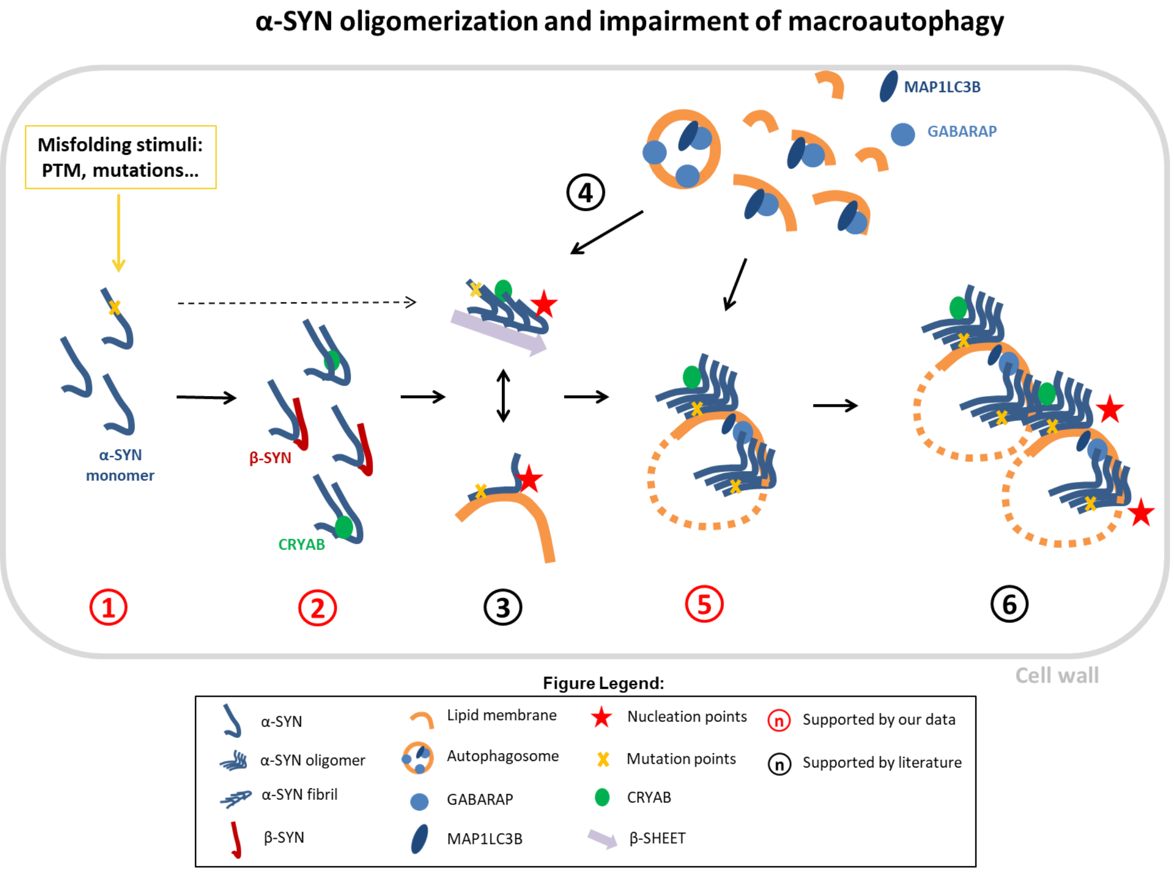


**Figure S11. Proposed model of early α-Syn aggregation and Lewy Body formation.**

**1-** Mutations and PTMs of the α-SYN molecule enhance its aggregation propensity. **2-** β-SYN and αβ-crystallin are chaperones of α-SYN that recognize early partially folded intermediates/aggregates, however faster-aggregating forms of α-SYN 'escape' proteostasis (dashed arrow). **3-** Membrane-bound oligomers and β-sheet fibrils coexist during the aggregation cascade of α-SYN and constitute nucleation points for further conversion and/or elongation. **4-** α-SYN aggregates co-diffuse in vitro with ATG8 proteins. **5 –** Co-diffusion of α-SYN oligomers with ATG8 proteins leads to growth of aggregates. **6-** Further growth and recruitment of aggregates potentially impairs autophagosome assembly. This supplementary figure refers to **Figure 6**.

### SUPPLEMENTARY APPENDICES

#### APPENDIX I: Cell-Free Methods to Measure Protein-Protein Interactions.

**Alphascreen assay:**

Direct interactions between monomeric N-terminal mCherry α-SYN and the N-GFP Lewy Body proteins were assessed using AlphaScreen. AlphaScreen is a nanobead assay, where the two interacting proteins bring a "donor" bead and an "acceptor" bead in close proximity. The donor bead is coated with streptavidin, which binds biotin-coupled GFP nanotrap. The GFP nanotrap recruits the GFP-tagged protein (LB protein). The acceptor bead is coated with anti-myc antibody that binds to the mCherry-myc tag of α-SYN. The donor bead contains phtalocyanine, a photosensitizer that converts ambient oxygen to an excited state upon illumination at 680 nm. The resulting singlet oxygen has a half life of 4µs, in which it can diffuse ~200 nm in solution, before returning to its triplet unexcited state. If the two proteins (N-Cherry α-SYN and N-GFP LB protein) interact, the beads are brought within that distance and the singlet oxygen reacts with thioxene derivatives in the acceptor bead, leading to emission of light at 520-620 nm. The technique has high sensitivity, thanks to the local increase of the concentration of the proteins at the surface of the beads, allowing the formation of low affinity complexes. However, AlphaScreen signal is dependent on the concentration of the protein: at optimal protein-to-bead ratios (when the beads are fully coated), a maximum signal is detected as shown by a typical 'hook effect' in the quantified luminescence emitted (**Figure 2-B**).
In this study we take advantage of another aspect of the system. We performed 4 serial dilutions of the samples containing the co-expressed proteins to find the optimal proteins-to-beads ratio. Typically, optimal coating of the beads is achieved when we dilute the LTE by 3-4 orders of magnitude. At these (lower) concentrations, dissociation rates are high and even aggregation-prone proteins will be largely monomeric. Therefore, the presence of few aggregates should not overwhelm the system and interactions between aggregates will have the same influence as interactions between two monomers.

**Two-color Coincidence Detection (TCCD) to study PPIs:**

Two-color coincidence detection (TCCD) refers to the simultaneous excitation of two fluorophores by two independent lasers and detection of the fluorescence emitted as a result. TCCD relies on the diffusion of a dual-labelled complex (such as an heterogenous aggregate of two differently tagged species) through the confocal volume, which gives rise to coincident bursts of fluorescence on both channels. This approach can be used to sensitively detect the presence of dual-labelled molecules, even in the presence of a large excess of singly labelled molecules, as are typically our cell-free expressed samples of fluorescent proteins. It can also be used to determine the stoichiometry of the complex (hence, of the interaction).
The identification of fluorescent events in TCCD is achieved by counting the number of photons emitted at a set interval time $\tau$. First, however, one must define what is to be considered an 'event'. This is achieved by establishing thresholds, above which bursts are recorded. The presence of two events on both channels at the same bin-time (selected by the operator) is read as a coincident event. The simple binning-threshold method for event selection functions extremely well providing that signal to noise ratio is sufficiently high, and the interval time is longer than the diffusion time of the molecule across the probe volume.
TCCD is now being applied to increasingly complex systems, bringing about issues in the detection of fluorescent events. Systems where the expression level of proteins is difficult to control or where background levels are too high may affect the threshold selection. Another factor we had to overcome in this study is the occurrence of chance coincident events, i.e. when two species tagged with different colours enter the confocal volume at the same time by chance. Ideally, chance coincident events should represent only a fraction of the total number of events, but sometimes chance for co-diffusion can increase. This is particularly critical in our studies of interactions with α-SYN fibrils, which were performed at single molecule concentrations, where dissociation rates can be higher.
To tackle these limitations the choice of threshold is critical, and here we followed an optimized methodology published by Clarke and colleagues[11], as described next.

Association Quotient (Q)
Choosing optimal thresholds to define an 'event' is a critical step in burst detection: the threshold should be high enough that exceeds the background level and minimizes detection of chance co-diffusions; it should also be low enough that ensures collection of enough fluorescent events.
Correction for chance can be incorporated into the calculation of an association quotient, Q, that defines a fraction of coincident events from the total number of events. Clarke *et al*[11] have demonstrated recently that appropriate thresholds can be calculated automatically by plotting the population of Q values as a function of the thresholds and finding the maximum Q values. According to the authors, Q is defined as:

$$Q = \frac{(C - E)}{(A + B - (C - E))} \qquad (1)$$

Where *A* and *B* are the events in the two channels, the observed rate of coincident events is C and the estimated rate of events that occur by chance is given by *E*, which in turn can be defined as:

$$E = AB\tau \qquad (2)$$

Where A and B are the total rates of events in the two channels and $\tau$ is the interval time in seconds. In **Figure 3-A-iii** we show a simple representation of a Q value calculated for a control co-expression of C-GFP-tagged G51D α-SYN with C-Cherry-tagged G51D. We used the approach described in the manuscript mentioned above[11] to calculate, for each time trace of co-expressed m-Cherry- and GFP-tagged proteins the optimal threshold, as shown in **Figure S3**.
In sum, by plotting thresholds against each corresponding Q value, we obtain a color-field graph such as represented in Figure S3, and we are then able to define the optimal threshold as the lower threshold that gives the maximum Q value. This is represented in our example in **Figure S3 (insets B and D)** as the darker warm colours and in the raw trace (**insets A and C**) as the blue lines. Below this threshold, the background is too high, increasing the chance ($E$) for non-interacting partners to co-diffuse (therefore decreasing $Q$ in equation 5); on the other hand, above this threshold we sample a number of events that is too low for a representative analysis. In the color-field maps above, we see high Q values at the top of the graph as well, owing to the very few events that are detected at such high levels, which will mostly be picked as coincident; hence we automatically select the lower threshold. This example shows the importance of threshold selection in our work, as we performed hundreds of coexpressions of proteins. Therefore, a fast, expedite and automated way of calculating association quotients in a high throughput manner was required.
By applying this methodology to all our co-expression traces we obtained consistent Q values, as shown by relatively low dispersity of replicates in our results section.

Coincidence ratio for stoichiometry analysis
In this work we calculated stoichiometries of interactions for identified binders of α-SYN in order to determine the relative presence of each species in heterogenous aggregates. We collected the population of coincident events (using the same threshold selection method as described above) and calculated a coincidence ratio $C$ that gives the average fraction of mCherry fluorophores in the aggregate, as defined by the formula below:

$$C = \frac{I_{GFP}}{(I_{Cherry} + I_{GFP})} \qquad (3)$$

Where $I_{Cherry}$ and $I_{GFP}$ are the threshold-subtracted intensity in the Cherry and GFP channels, respectively. It should be noted that before applying this fraction for each event, the intensities of the GFP and Cherry bursts were corrected for background and leakage (6% leakage of the GFP intensity into the Cherry channel). By applying equation 3 to our population of coincident events we obtain a number between 0 (absence of GFP fluorescence) and 1 (absence of Cherry fluorescence). Histograms of single-molecule coincidence were obtained by measuring >400 events per interaction.

#### APPENDIX II: Investigating the presence of functional LIR motifs in Lewy Bodies.

If we consider the hypothesis that pathological α-SYN aggregates could overwhelm autophagic fluxes by binding/clustering to autophagosomes, it is then also worth investigating which other proteins in the Lewy Bodies bind to these structures and potentially increase convolution. We have scanned the list of Lewy Body components for the presence of LIR motifs that are required for binding to autophagosomal (ATG8) proteins, using a bioinformatical consensus sequence search engine[58] (**Figure S12**). Kalvari and colleagues[48] have recently refined the LIR motif to an extended LIR (xLIR) that represents functional motifs (experimentally validated) with higher accuracy than the canonical LIR (cLIR). Accuracy for functional LIR detection can be further increased when used in conjunction with a position-specific-scoring-matrix (PSSM) that analyses the probabilistic multialignment fit. In addition, LIR motifs that overlap with an intrinsically disordered region, as detected by ANCHOR software are more likely to represent functional motifs. This is because LIR's are generally extended sequences that exhibit characteristics of conformational switches, i.e. they fold upon binding to autophagosomes. Using this method we have detected with high accuracy at least three LIR-containing proteins in our list of LB proteins: gelsolin (GSN), neighbour to BRCA1 (NBR1) and NF-kappa-β inhibitor alpha (NFKBIA). While NBR1 is known to be involved in delivery of protein aggregates to autophagosomes for degradation, GSN and NFKBIA have not been identified before as binders of ATG8 proteins and could be important regulators of autophagy.

Coincidently, specific α-SYN aggregates have been shown to co-diffuse with all these proteins in our PPI screen. Interactions between α-SYN aggregates and these LB components could, in principle, lead to generalized loss of function and increase the number of species that become trapped in insoluble inclusions, potentially amplifying the collapse into the non-functional protein clumps that are Lewy Bodies.

#### SUPPLEMENTARY FIGURE 12

*cLIR:* [DE][DEST][**WFY**][DELIV]X[**ILV**]

*xLIR:* [ADEFGLPRSK][DEGMSTV][**WFY**][DEILQTV][ADEFHIKLMPSTV][**ILV**]

| GENE | ID | Family | Motif (i) | Start | End | Pattern | PSSM (ii) | LIR in IDR (iii) | cLIR + IDR | xLIR+IDR+P>13 | xLIR+IDR\|P>13 | OVERALL |
|---|---|---|---|---|---|---|---|---|---|---|---|---|
| BTRC | Q9Y297 | Ubiquitin | xLIR | 529 | 534 | K V W D L V | 13 | No | | | | |
| | | | xLIR | 564 | 569 | D E F Q I V | 14 | No | | | | |
| CAMK2B | Q13554 | Kinase | xLIR | 88 | 93 | L V F D L V | 5 | No | | | | |
| | | | xLIR | 236 | 241 | P E W D T V | 19 | No | | | | |
| CCNB1 | P14635 | Mitosis | xLIR | 332 | 337 | L D Y D M V | 16 | No | | | | |
| CSNK2A1 | P68400 | Kinase | xLIR | 10 | 15 | R V Y T D V | 7 | No | | | | |
| | | | xLIR | 37 | 42 | D D Y Q L V | 17 | No | | | | |
| | | | xLIR | 52 | 57 | E V F E A I | 8 | No | | | | |
| | | | xLIR | 111 | 116 | L V F E H V | 5 | No | | | | |
| CSNK2A2 | P19784 | Kinase | xLIR | 38 | 43 | D D Y Q L V | 17 | No | | | | |
| | | | xLIR | 53 | 58 | E V F E A I | 8 | No | | | | |
| GSN | P06396 | Actin binding | xLIR/cLIR | 472 | 477 | D S Y I I L | 15 | Yes | | | | * |
| | | | cLIR | 702 | 707 | D T W D Q V | 19 | Yes | | | | |
| HSP90AB | P08238 | Chaperone | xLIR | 506 | 511 | R G F E V V | 12 | No | | | | |
| HSPD1 | P10809 | Chaperone | xLIR | 126 | 131 | E G F E K I | 12 | No | | | | |
| HSPH1 | Q92598 | Chaperone | xLIR | 87 | 92 | L S Y D L V | 10 | No | | | | |
| | | | xLIR | 714 | 719 | K M F E E L | 12 | No | | | | |
| NBR1 | Q14596 | autophagosome | xLIR | 561 | 566 | L S F E L L | 10 | Yes | | | | |
| | | | xLIR/cLIR | 730 | 735 | E D Y I I I | 17 | Yes | | | | * |
| NFKB1A | P25963 | transcription factor | xLIR/cLIR | 75 | 80 | D S F L H L | 13 | Yes | | | | * |
| PIN1 | Q13526 | isomerase | xLIR/cLIR | 101 | 106 | E D F E S L | 16 | No | | | | |
| PLCD1 | P51178 | lipase | xLIR | 5 | 10 | R D F L T L | 16 | No | | | | |
| PSMC6 | P62333 | proteasome | xLIR | 265 | 270 | D G F D T L | 15 | No | | | | |
| SLC18A2 | Q05940 | transport | xLIR | 73 | 78 | D S F Q S I | 13 | No | | | | |
| SNCAIP | Q9Y6H5 | interacting partner | xLIR | 4 | 9 | P E Y L D L | 12 | No | | | | |
| | | | xLIR | 166 | 171 | A S F T K V | 9 | No | | | | |
| SYP | P08247 | membrane association | xLIR | 26 | 31 | L G F V K V | 9 | No | | | | |
| | | | xLIR | 123 | 128 | A T Y I F L | 12 | No | | | | |
| SYT11 | Q9BT88 | membrane association | xLIR | 299 | 304 | L S Y Q P V | 9 | No | | | | |
| TGM2 | P21980 | enzyme | xLIR | 352 | 357 | E G W Q A L | 18 | No | | | | |
| TOR1A | O14656 | enzyme | xLIR | 126 | 131 | S D Y V H L | 17 | No | | | | |
| | | | xLIR | 201 | 206 | A M F I F L | 11 | No | | | | |
| UBA1 | P22314 | ubiquitin | xLIR | 157 | 162 | S G F Q V V | 11 | No | | | | |
| | | | cLIR | 650 | 655 | D E F E G L | 13 | No | | | | |
| VPS26B | Q4G0F5 | retromer | xLIR | 107 | 112 | F E F T H V | 11 | No | | | | |
| MAP1B | P46821 | cytoskeleton | xLIR | 252 | 257 | G G F L K L | 10 | No | | | | |
| | | | xLIR | 838 | 843 | K D F E E L | 16 | Yes | | | | |
| | | | xLIR | 2355 | 2360 | P V Y L D L | 7 | Yes | | | | |
| SQSTM1 | Q13501 | autophagosome | xLIR | 336 | 341 | D D W T H L | 24 | Yes | | | | |
| | | | xLIR | 296 | 301 | D V F D D I | 10 | Yes | | | | |
| MAP2 | P11137 | cytoskeleton | xLIR | 1243 | 1248 | G E Y D K L | 13 | Yes | | | | |
| | | | xLIR/cLIR | 1289 | 1294 | D D F I T V | 17 | Yes | | | | |

| | cLIR + IDR | xLIR+IDR+P>13 | xLIR+IDR\|P>13 |
|---|---|---|---|
| **Accuracy (%)** | 57.4 | 70.2 | 78.7 |
| **Sensitivity (%)** | 29.6 | 55.6 | **96.3** |
| **Specificity (%)** | **95** | **90** | 55 |
| **Balanced accuracy (%)** | 62.3 | 72.8 | 75.7 |

**Figure S12. Searching for functional LIR motifs in the LBs. (A)** Canonical and extended LIR motifs (cLIR and xLIR respectively) were detected using a LIR search engine available online and compared against known LIR containing proteins (MAP1B, SQSTM1 and MAP2, bottom rows). This search engine detects the consensus sequences surrounding a core sequence of aromatic (W/F/Y) and hydrophobic aminoacids (I/L/V) – highlighted in yellow. Functional LIR motifs were selected using 3 factors (highlighted in red): **i)** presence of cLIR/xLIR; **ii)** overlap (more than 3 residues) with intrinsically disordered region (IDR), as detected by ANCHOR software; and **iii)** position specific scoring matrix (PSSM) >13. Using different combinations of these 3 factors (cLIR+A, xLIR+IDR+PSSM>13 and xLIR+IDR or PSSM>13), three functional LIR containing proteins were detected with high confidence (green in 'OVERALL' column), based on selectivity and specificity of each test: GSN, NBR1 and NFKBIA. Other proteins that could contain functional LIR motifs are represented in blue (however this is unclear due to low specificity). **(B), (C)** Proteins containing functional LIR motifs co-aggregated with α-SYN, which could have deleterious consequences for the overall autophagic flux.
